# Modular and redundant genomic architecture underlies combinatorial mechanism of speciation and adaptive radiation

**DOI:** 10.1101/2025.07.07.663194

**Authors:** Pooja Singh, Heidi E.L. Lischer, Kassandra Ford, Ehsan P. Ahi, Marcel P. Haesler, Salome Mwaiko, Joana I. Meier, David Marques, Remy Bruggman, Mary Kishe, Ole Seehausen

## Abstract

Hybridisation can fuel rapid adaptive radiation, but how it enables the formation of phenotypically highly dimensional species-rich radiations remains unclear. We investigated this by analysing genotype-phenotype associations for 14 ecological (trophic, body patterns) and mating traits (nuptial colour) across 107 species of Lake Victoria cichlid fishes. We find weak trait covariance across the radiation, with many different trait combinations constituting different species. Across the radiation, polygenic, redundant, and lowly pleiotropic genomic architectures of hybrid origin underlie the repeated evolution of key traits. Such independent genomic modules can be reshuffled and recombined like Lego bricks, generating diverse trait combinations from a finite number of elements. During speciation, dispersed oligogenic trait modules become coupled through long-range linkage disequilibrium. We propose that this genomic and phenotypic modularity emerged from repeated cycles of past hybridisation, enabling superfast adaptive radiation through combinatorial speciation.

## Main Text

Adaptive radiation, where an ancestral lineage diversifies quickly into many eco-morphologically diverse species that can also quickly coexist in species rich communities (non-geographical radiation), may have been an important source of biodiversity in key moments of the 3.5-billion-year history of Life on Earth (*1–3*). Its occurrence is highly punctuated in space, time, and phylogeny, and despite its likely importance, knowledge of the biological factors that trigger, facilitate, and constrain this process is incomplete (*4*, *5*). It is widely accepted that exceptional ecological opportunity is a prerequisite (*6*), but only some of many possible lineages seem to ever have responded to exceptional ecological opportunity with adaptive radiations and even fewer have done so rapidly and repeatedly. From studying some of the geologically youngest radiations, it is apparent that the process can sometimes unfold in just a few thousand years (*7–9*).This is consistent with the fossil evidence for long periods of relative stasis, punctuated by bursts of diversification in the history of life(*10*, *11*). Such species radiations, with many speciation events in succession are far too fast for intrinsic reproductive isolation to have emerged before species assemble into rich sympatric communities. But they are also far too young for adaptive variation to have arisen through mutations and tend to be characterised by the coincidence of hybrid ancestry and/or recurrent hybridization and simultaneous evolution of ecological and mating traits, as was predicted by Anderson & Stebbins 70 years ago (*10*). While some inroads have been made linking rapid evolution to the availability of ancient admixture variation in plants and animals (*9*, *12–17*), we still understand little about the genetic basis that not only allows key traits to evolve rapidly, but that permits the rapid generation of many different highly dimensional phenotypes that are associated with many speciation events in short succession in the absence of geographical isolation.

Research has illuminated two prevalent genetic architectures underlying ecological speciation without geographical isolation. These architectures are either (1) simple and consist of a few genomic ‘islands’ of differentiation made up of either clustered small effect alleles physically linked into a supergene or single loci of large effect (*18–21*) or (2) more complex and involve genome-wide and/or polygenic patterns of differentiation (*22–33*). Each of these architectures imposes unique constraints on speciation in the face of gene flow (*23*, *34*). Under the scenario of simple concentrated trait architectures, phenotypic differentiation may be maintained despite some gene flow because total selection will be distributed over one or a few regions in the genome and may hence be strong enough to counter gene flow at those loci and also because large effects of single loci cannot be broken down by recombination (*35*, *36*). However, speciation is improbable in the first place unless if the locus under divergent selection also causes reproductive isolation as a by-product, e.g. in the form of ‘magic traits’ (*37–39*) or through physical linkage to loci causing reproductive isolation (*40–42*). Indeed, adaptive polymorphism within species have been attributed to such genetic architectures and have proven not to be associated with speciation, including in cichlid fish (*43*, *44*). When trait architectures are complex such that expression of divergent trait values requires many small to medium effect alleles distributed over multiple chromosomes, it is more likely that some of these loci are either physically linked to genes affecting mating compatibility or affect the latter through pleiotropy. If not, the strength of divergent selection per locus will likely be insufficient to counteract the homogenising effects of gene flow at any one locus, making speciation without geographic isolation unlikely (*34*, *45*). Theory suggests that the latter constraint can be overcome either through the gradual build-up of weak polygenic Linkage Disequilibrium (LD) at many loci across the genome over millions of years, followed by rapid transition to two species around genomic ‘tipping points’(*46*) or through hybridisation introducing many divergent haplotypes with multiple linked SNPs at once (*47*, *48*). Neither of these theories have robustly been tested empirically in any adaptive radiation. Furthermore, most theoretical speciation-with-gene flow models were built to explain singular speciation events where the processes of accumulation of variation and building of LD restart after every speciation event. Such models cannot easily be applied to adaptive radiations with tens or hundreds of speciation events that occurred within ten or hundred thousand years (but see (*47*, *49*)).

Adaptive radiations that produce many sympatric species, whether in cichlids, Darwin’s Finches or Hawaiian Honeycreepers, are associated with high dimensionality of phenotypic and ecological variation even early on in the process. Hence, selection was most likely diverse and multifarious. Few theoretical and even fewer empirical studies have considered the genetic architecture that allows many successive or simultaneous speciation events when the interaction of multiple environmental factors impose divergent selection pressure on multiple traits and trait combinations, and when there is not sufficient time between successive speciation events for recovery of genetic variation through mutation (*48*, *50*). Here, we addressed this problem in the fastest and most species-rich cichlid fish adaptive radiation known by conducting radiation-wide genotype-phenotype association analyses of hundreds of individuals that represent 107 species and 14 ecological and mate-choice traits (Fig. 1).

**Fig. 1.**
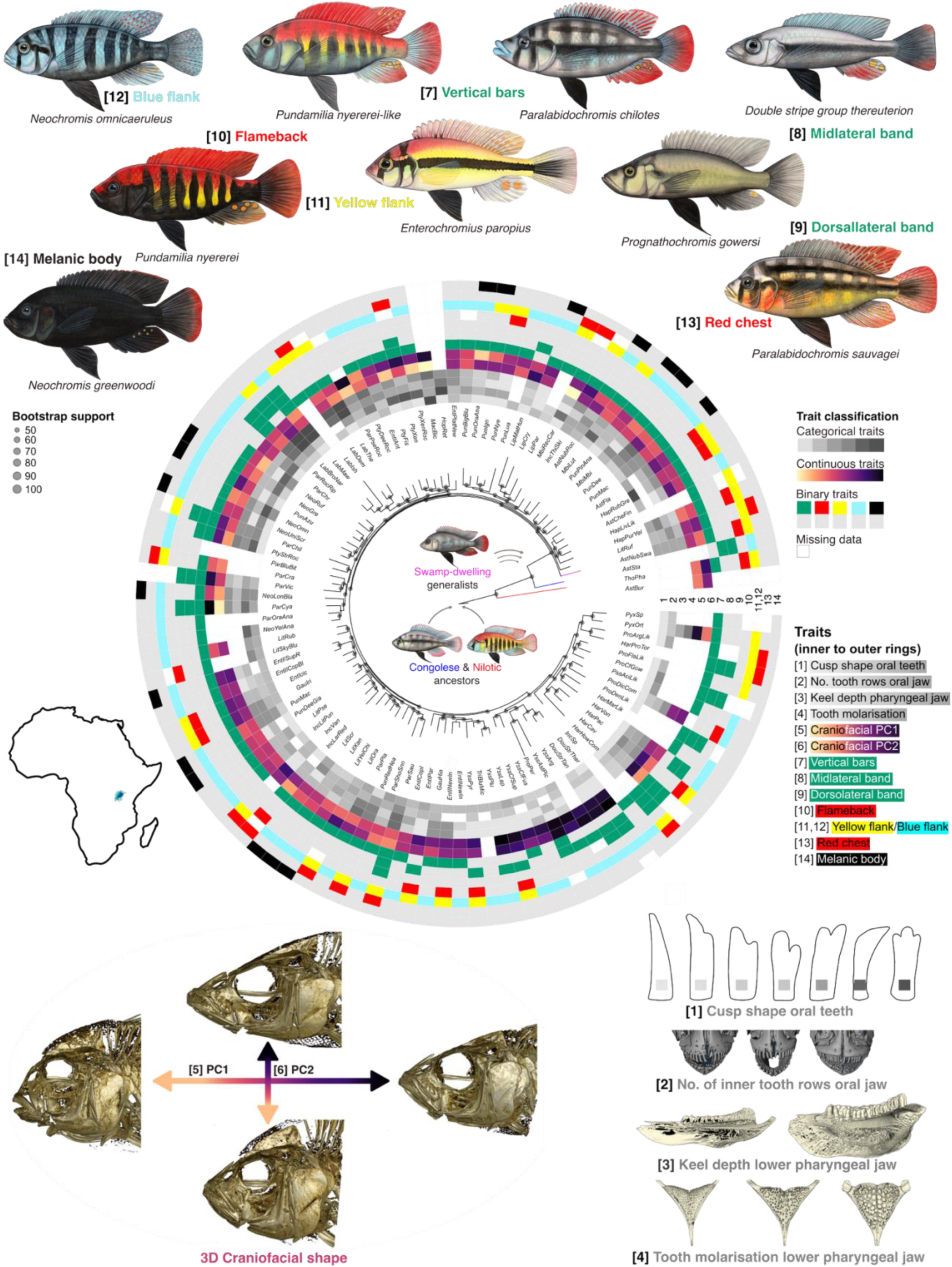
Overview of ecological and mating trait evolution in the Lake Victoria cichlid adaptive radiation. The Lake Victoria cichlid radiation encompasses ∼500 known species (107 mostly sympatric species shown in the phylogeny) that represent unique phenotypic and genetic mosaics of recurrent ecomorphologies, melanic stripe patterns, and male nuptial colour motifs (illustrated with species shown), assembled in <16,000 years through combinatorial mechanisms (*51*). Distribution of 14 core traits used in this study are shown in 13 rows (traits 11 and 12 are mutually exclusive traits and thus plotted in the same row)around the genome-wide SNP phylogeny (constructed using *P. nyererei* v3 reference genome here) in such a way that each column ray around the circle tree represents the unique set of trait combinations present in the species at the corresponding tip of the tree. Ecomorphological traits are numbered from 1 – 6: cusp shape of outer row frontal teeth [1] and number of inner tooth rows in oral jaws [2], lower pharyngeal jaw keel depth [3] and molarisation of lower pharyngeal jaw teeth [4], as well are 3-dimensional (3D) craniofacial shape PC1 [5] and PC2 [6]. Colour gradients of PC1 and PC2 arrows correspond to the PC value heatmap as illustrated around the phylogeny. Melanic stripe patterning traits are numbered from 7 – 9: vertical bars [7], midlateral band [8] and dorsolateral band [9], where the latter only occurs in combination with [8]. Male nuptial colour motifs are numbered from 10 – 13: flameback [10], yellow or blue flank [11,12], red chest [13], and melanic body [14]. Traits 1 – 4 are categorical (see Fig. S1 for all categories), traits 5 and 6 are continuous, and traits 7 – 14 are coded as binary (presence/absence). Node bootstrap support as circles of increasing sizes. Key ancestral admixture events in the ancestry of the radiation from (*9*) are shown. Distribution of ecological and mating traits across the phylogeny suggests that trait modules can largely recombine freely into different combinations in species.

### Low covariance among phenotypic modules of adaptive radiation

If admixture variation from the hybrid swarm origin of the radiation was important to permitting exceptionally fast onset of speciation and adaptive radiation when the new lake formed, and if continued syngameon conditions in the emerging radiation were key for maintaining the momentum through many successive speciation events, we expect a radiation-wide genome panel to uncover the recycled component of the genetic architecture of key traits associated with repeated use of the same alleles for the same traits across the radiation, albeit in different trait combinations in different species (*52*). We explored this expectation with a Radiation-Wide genotype-phenotype Association Study (RWAS) after generating a long-read improved reference genome assembly and Iso-seq based gene annotation (Table S1). To this end we conducted the most comprehensive phenotyping in any cichlid adaptive radiation to date and analysed genomes of 324 individuals from 107 different species (File S1). Unlike GWAS on a few species or a hybrid population, the RWAS approach is not designed to capture genetic variants that contribute to a trait in just one or two species but is designed to capture variants only if they affect the same trait in several species, which would be a prediction of the combinatorial hypothesis for adaptive radiation (*51*). We focused on 14 key traits involved in ecological adaptation and mating behaviour (Fig. 1) that make up three complementary trait complexes: (1) ecomorphology - we quantified craniofacial shape using 3D geometric morphometric from high resolution CT scans and summarised it as continuous variation along major axes of a Principal Component Analysis based on 3D geometric morphometrics from high resolution CT scans (Fig. S1); we categorised shape and arrangement of teeth in the oral jaws, as well as pharyngeal jaw shape and tooth molarisation into five to seven categories (Fig. S2); (2) melanic stripe patterns - presence/absence of vertical bars on the flank, a midlateral band, and a dorsolateral band (only occurs in conjunction with midlateral band) (*53*); (3) male nuptial colour motifs - we scored presence/absence of all known recurrent Lake Victoria cichlid male nuptial colour motifs (*54*, *55*): blue flank versus yellow flank, flameback (red-dorsum), red chest (red-ventrum), and melanic body (Fig. 1). Craniofacial shape, pharyngeal and oral jaw shape and dentition are all correlated with trophic ecology (*56*, *57*), melanic stripe patterns are thought to play a role in adaptation to different habitats and shoaling versus territorial behaviour (*53*) and male nuptial colouration is of critical importance in female mate choice, male territoriality and thus, species persistence in sympatry (*58–61*). Together, these 14 traits capture major aspects of the interspecific variation that characterises the Lake Victoria cichlid radiation (*56*, *57*).

Our phenotyping revealed that across the radiation these 14 traits may be quite freely recombined with each other, allowing a very large number of unique species phenotypes to be generated just from this trait set (see Fig. 1). A few pairs of correlated traits exist, but most traits appear to evolve independently of each other (mean absolute Spearman’s ρ correlation coefficient 0.27; Fig. 2a). Noteworthy positive correlations exist between: yellow flank and flameback, whereas blue flank is exclusive of yellow flank and negatively correlated (but not exclusive) with flameback; midlateral and dorsolateral band, where the latter does not occur without the former; lower pharyngeal jaw keel depth and the extend of tooth molarisation. Noteworthy negative correlations exist between craniofacial shape PC1 (elongation of head and snout and upwards/downwards orientation of mouth) and number of tooth rows in the oral jaws (a high number of tooth rows is seen only in fish with downwards opening mouths); and between midlateral band and vertical bars. All of these recover well-known between-species trait associations (*56*, *62*, *63*). The fact that most other traits are only mildly correlated across the radiation is noteworthy because highly correlated traits would constrain diversification to a limited number of unique phenotypes and species, while uncorrelated traits can be freely combined into many unique species in a modular fashion (*64*).

**Fig. 2.**
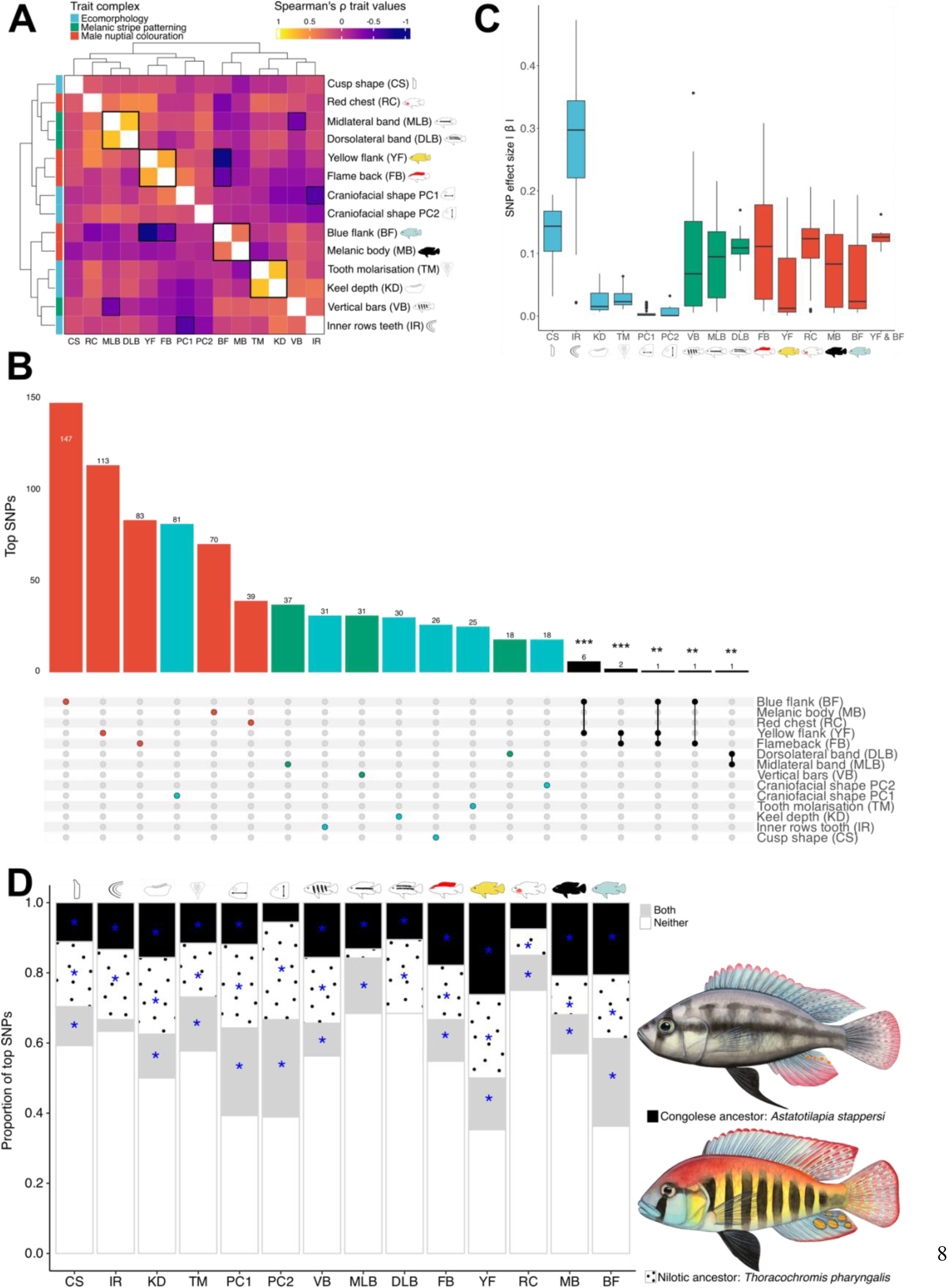
Radiation-wide association analyses (RWAS) reveal polygenic trait architectures and their admixture history. (**a**) Spearman’s ρ correlation coefficients between ecological and mating traits across all species in this study. Some pairs of traits show co-variation across species (highlighted with black squares; see text for details), but most traits do not covary strongly. (**b**) The repeatedly used component of genetic architecture of all traits is polygenic as demonstrated by the number of RWAS top SNPs significantly associated with each trait (top 99.99^th^ percentile & PIP > 0.01 for binary/categorical traits and top 99.9^th^ percentile & PIP > 0.01 for continuous traits; from here on denoted top SNPs). Little overlap of top SNPs among traits is interpreted as low pleiotropy among these traits (Hypergeometric test *p* values denote significant overlaps: *p* < 0.001***, *p* < 0.01**). (**c**) The absolute effect size (|β|) distribution of top SNPs for each trait shows some medium to larger effect SNPs and many small effect SNPs. (**d**) Most trait-associated top SNPs are also found in the closest living relatives of the Congolese and/or Nilotic ancestors of the Lake Victoria region superflock. Their origins hence predate not only the Lake Victoria cichlid radiation but the entire Lake Victoria Region species flock which includes radiations in Lakes Edward, Albert, Kivu and smaller satellite lakes. For each trait category we tested whether ancestral lineage SNPs are overrepresented among the trait-associated SNPs in the radiation (observed value exceeds 99th percentile of 100 iterations of randomly sampled SNPs across the genome; significantly enriched trait categories are indicated with an asterisk (*p* < 0.05*).

### Genomic architecture of ecological and mating traits is modular

Our RWAS across 107 species revealed that the recurrently used components of genetic architectures of all 14 key traits were polygenic at the level of the radiation, varying in their complexity ranging from 18 to 147 significantly associated SNPs hereon denoted top SNPs per trait (top 99.99^th^ percentile & PIP > 0.01 for binary/categorical traits and top 99.9^th^ percentile & PIP > 0.01 for continuous traits (*65*)) (Fig. 2b). These top SNPs were dispersed across the genome for all traits (Fig. 3a, Fig. S3). Craniofacial shape and male nuptial colouration traits had the greatest number of top SNPs, followed by melanic stripe patterns, dentition and pharyngeal jaw morphology. With the RWAS approach it is likely that we detect fewer top SNPs for more complex and redundant trait architectures, with many alleles of small effect, different subsets of which could be used in each species (i.e. less recurrent use across the radiation). While traits with genetic architectures that are simple and repeatedly used across many species are statistically more likely to be detected. This is probably why we find more top SNPs for male nuptial colour motifs than for most ecomorphological traits (Fig. 2b). This is especially true for craniofacial shape, presumably a set of complex traits in itself that is summarised in dimension-reduced PC axes (Fig. 1, Fig. S1). The effect sizes of top SNPs varied within and between ecological and mating trait complexes (Fig. 2c). As expected, the presumably more quantitative traits such as craniofacial shape PCs and pharyngeal keel depth were entirely determined by alleles of small effect, while more categorical dentition traits such as the number of inner tooth rows in the oral jaw and cusp shape of outer row oral teeth had some of the largest effect sizes. Interestingly, we found that the pharyngeal jaw tooth molarisation was entirely determined by alleles of small effect, which is a sharp contrast to the simple architecture of this trait in the polymorphic central American cichlid (*43*). Melanic stripe patterns and male nuptial colour motifs were characterised by small to medium effect alleles (Fig. 2b). A detailed discussion of candidate genes and GO enrichment (Fig. S4) for all traits can be found in the supplementary material. Next, we applied random forest machine learning to evaluate the predictive accuracy of the top SNPs for the eight binary body colour and patterning traits. Models were implemented within a five-fold cross-validation scheme, where in each fold 80% of individuals were used for training and 20% were held out for testing.

**Fig. 3.**
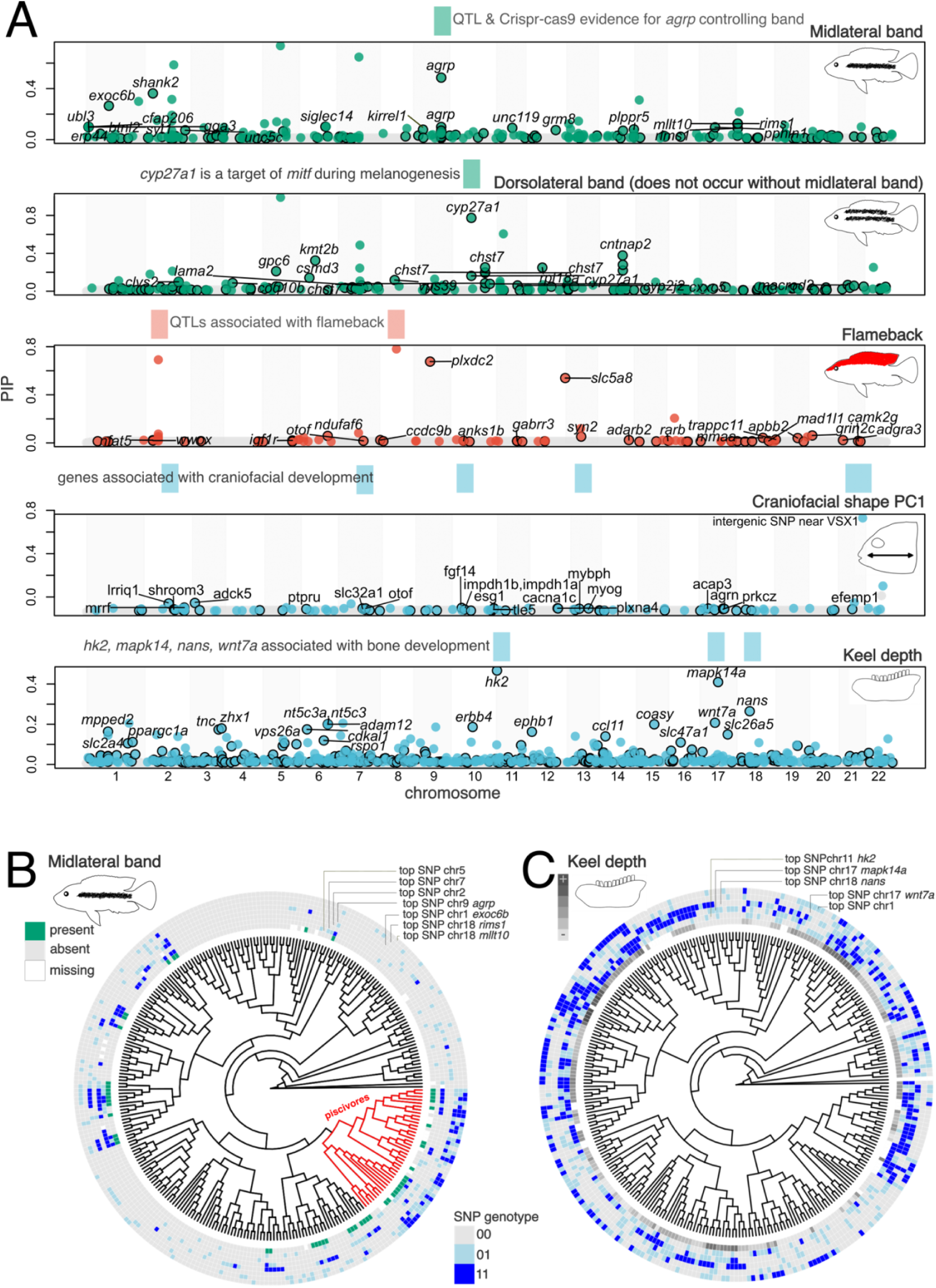
Genomically dispersed and redundant trait architectures in Lake Victoria cichlids. **(a)** RWAS top SNPs repeatedly associated with key adaptive radiation and speciation traits in the Lake Victoria radiation are dispersed across the genome. Significant top SNPs are highlighted as coloured circles and non-significant SNPs are denoted by grey circles (barely visible at the bottom of the Y axes). Colour of circles are categorised by trait complexes, colour coded as in Fig. 2: blue – ecomorphology, green-melanic stripe pattern, red-male nuptial colour. Coloured circles with black outline are SNPs located in genes and coloured circles without black outline are SNPs located in intergenic regions. For each trait the top 20 genic SNPs are annotated with gene names - full list can be found in File S1. Functional evidence linking candidate genes to traits from previous QTL mapping and gene knock-out studies (cited in main and supplementary text) for each trait is illustrated above each Manhattan plot (see Fig. S3 for all 14 key traits). (**b, c**) Two examples of evolutionary genetic redundancy in key traits: midlateral and lower pharyngeal jaw keel depth repeatedly evolved repeatedly across the LV radiation, but different top SNPs/genes are associated with the same trait in different species and/or sub-clades in the radiation. Genetic redundancy plots for the other 12 traits are presented in Fig. S8.

Across traits, SNPs with higher PIP score were consistently associated with greater importance in phenotype prediction models. Specifically, we observed significant positive Pearson correlations between SNP PIP scores and their Mean Decrease in Accuracy (MDA) values from the Random Forest models for all traits except Dorsolateral band (Fig. S5, Fig. S6). Correlation coefficients ranged from 0.31 to 0.78 (*p* < 0.05 for each trait), indicating substantial concordance between RWAS-derived association strength and machine learning–based predictive importance (Fig. S5).

Surprisingly, there was little overlap of top SNPs between traits (Fig. 2b) and this overlap did not increase substantially even when we considered overlap at the level of genes containing top SNPs (Fig. S7). This suggests low pleiotropy across the ecological and mating traits that are associated with the radiation, such that variation in these traits can be largely independently inherited. Such modular architectures may have been important for allowing many different trait combinations to evolve in very little time, permitting the differentiation of this lineage into hundreds of different and phenotypically unique species. Those cases of pleiotropy that we did find are consistent with what is known from patterns of co-variation across species in nature and in species crosses in the lab, particularly among nuptial colour motifs and among stripe pattern elements (Fig. 2a). For instance, the largest overlap were six top SNPs shared between Yellow flank (YF) and Blue flank (BF) (hypergeometric test *p-value* < 0.001), which are divergent and mutually exclusive male nuptial colours in the iconic *Pundamilia* sister species (Fig. 2b) and inversely correlated across the radiation (Spearman ρ = -0.8; Fig. 2a). These six SNPs that had opposite effect sizes on these traits (+β on BF vs -β on YF) mapped to *clul1*, a cone photoreceptor gene that is differentially expressed in light and dark-adapted retinas (*66*), and *pdfgrb*, a vertebrate colouration gene(*67*). Male nuptial colouration and the ability to visually detect it are expected to be correlated (*68*). Interestingly, we found *clul1* expressed in the eye transcriptomes of *P. pundamilia* that has blue flank male nuptial colour and blue-shifted colour perception to live in light-dark contrasting rock crevices in very shallow water (*68*). This gene was not expressed in the eye transcriptome of its sister species *P. nyererei* that lives in deeper waters, has flameback and yellow-flank male nuptial colour and red-shifted colour perception (*68*). It is possible that the top SNP which sits in the intron of *clul1* may disrupt its expression. Other relevant cases of pleiotropy at the SNP and/or gene level were between Midlateral band (MLB) and Dorsolateral band (DLB) – the latter is never expressed in Lake Victoria cichlids without the former; Vertical bars (VB) and MLB – the occurrence of which are negatively correlated; and the extent of lower pharyngeal jaw tooth molarisation and keel depth, which are strongly positively correlated (*56*). In summary, the few trait correlations that we observe (Fig. 2a) seem at least partly caused by pleiotropic gene action, but there is no evidence for pleiotropic gene action on other trait combinations.

### Trait modules are derived from ancestral hybridisation

Despite the extraordinarily young age of the Lake Victoria cichlid fish radiation, its species have acquired highly specialised trophic adaptations as well as a large array of nuptial colouration motifs and stripe patterns that can be combined with each other and with trophic morphology in many ways (*56*, *69*, *70*). To better understand the extent to which resuspension of ancient admixture variation through cycles of fusion and fission processes (*9*) may have been important in enabling this process, we checked how many RWAS top SNPs with statistical effects on craniofacial shape, dentition, pharyngeal jaw morphology, male nuptial colour motifs, our body stripe patterns could be traced back to the extant Congolese and Nilotic haplochromine species that are the closest known living relatives of those species that hybridised ∼150,000 years ago to give rise to the Lake Victoria region super-flock of radiations (*71*). Those same genetic variants must then also have been cycled through the fission and fusion round associated with the period of total desiccation of Lake Victoria and subsequent recolonisation by refugial populations 16,000 years ago (*9*). We found that up to ∼30% of the top RWAS alleles were unique to one of the ancestral lineages (Fig. 2d). Nilotic ancestor-derived variants had significantly higher than expected contributions to genetic variation underlying all traits except midlateral band (>99th percentile of 100 iterations of randomly sampled SNPs across the genome). Congolese ancestor-derived variants had significantly higher than expected contributions to genetic variation underlying all traits with the exception of red chest and craniofacial shape PC2 traits but including midlateral band. This is notable because the midlateral band trait is found in the closest living relative of the Congolese ancestors but not in those of the Nilotic ancestor (Fig. 2d fish drawings). Alleles of the *agouti-related protein* (*agrp*) in particular were contributed to the LV radiation from their Congolese ancestors. The category of top alleles that we could not trace to either of the ancestral lineages was not significantly overrepresented for any trait.

This suggests that the combination of ancestral genetic variants, combined into the ancestor of the radiation through interspecific hybridisation, was important for the trait diversification that characterizes this radiation.

It is likely that some of the alleles that were carried into the ancestral hybrid swarm have since been lost in the closest living relatives of the ancestors that we sequenced, and we will have missed others in our limited sampling of these lineages compared to the large number of Lake Victoria cichlids that we sequenced. Hence, 30-50% of trait-associated alleles derived from admixture is likely an underestimate. The low pleiotropy among traits that we found would also be consistent with alleles undergirding those traits being old, selection-tested variants that have been swapped repeatedly among lineages through hybridization.

### Genetic redundancy contributed to rapid adaptive radiation

Genetic redundancy is hypothesised to promote evolvability by providing multiple paths for similar evolutionary change (*72–75*). Genetic redundancy has rarely been considered in the context of adaptive radiation, but it can be a powerful mechanism to promote rapid repeated trait evolution during speciation. We found evidence of genetic redundancy in trait architectures of all of the 14 repeatedly evolving traits in the Lake Victoria cichlid radiation (Fig. 3b, Fig. S8). One noteworthy example was in the evolution of a midlateral band (Fig. 3b). We mapped the top SNPs for midlateral band (Fig. 3b) across the phylogeny and discovered that non-piscivorous species with midlateral band are almost always fixed for the alternative *agrp* allele (G/G) but piscivore lineages, many species of which also have a midlateral band, do not carry this allele (A/A) (Fig. 3b). The evolution of midlateral band may be driven by *exoc6b* in the piscivores, alleles of which segregate almost perfectly with the presence of the midlateral band in heterozygous (C/G) or homozygous state (G/G). *Exoc6b* plays a crucial role in the transfer of melanin from melanocytes to keratinocytes in *in vitro* experiments(*76*). We next asked if genetic redundancy for this trait (and likely others) was generated by the combination of ancestral gene pools into the hybrid swarm ancestor of the radiation. While we could map the *agrp* G allele to the Congolese ancestor that possess a midlateral band, but the *exoc6b* G allele could not be mapped to either the Nilotic or Congolese ancestor (File S1). This could be simply a consequence of small sample size of ancestral lineages or it could reflect *de novo* evolution. We found similar patterns of genetic redundancy among species with greater pharyngeal keel depths where *hk2/mapk14a* and *nans/wnt7a* appear to function redundantly, but also additively in some species (Fig. 3c). Alleles for *hk2/mapk14a* and *nans* were present in both ancestral lineages but *wnt7a* allele was contributed exclusively by the Congolese ancestor (File S1). Given the repeated cycles of hybridisation in the ancestry of the Lake Victoria cichlid radiation (*9*, *71*), it is likely that genetically redundant alleles for the same traits arrived from different ancestral lineages from across the African continent (Fig. 2d). Such redundant genetic architectures emerging from hybridisation may be a previously underappreciated factor that provides a larger pool of genetic variation for repeated evolution of traits. This would promote the simultaneous and rapid evolution of many different species with different combinations of a limited set of phenotypic traits in this adaptive radiation.

### Long-range linkage disequilibrium couples dispersed trait modules during combinatorial speciation

To test predictions of combinatorial speciation that we hypothesise to be responsible for the rapid emergence of several hundred distinct species in record time, we combined RWAS information on the genetic architecture and admixture history of recurrent allele use underlying major traits of adaptive radiation with estimates of the genomic landscape of speciation and species differentiation as inferred from genome wide divergence (F_ST_) among closely related sympatric species. A unique prediction of the hypothesis of adaptive radiation through combinatorial speciation (combinatorial adaptive radiation) is that each species pair in the radiation is characterised by unique patterns of linkage disequilibrium (LD) among different subsets of reused trait-associated alleles (*51*). Here we set out to perform the first test of this prediction in any adaptive radiation. We calculated F_ST_ landscapes and determined differentiation outlier loci (top 5% F_ST_) for 12 different sympatric pairs of closely related species in the Lake Victoria radiation. We then annotated F_ST_ outlier loci with top SNPs from our RWAS to identify in each species pair the subset of their ‘speciation loci’ (defined as loci displaying significantly stronger differentiation than the rest of the genome, in full sympatry; RWAS top SNPs that are also top 5% F_ST_) and tested for LD among these loci. Firstly, we found that the genetic architecture of traits of speciation was oligogenic (File S1) and involved fewer loci than the polygenic global architecture of these same traits across the radiation (Fig. 2, File S1). Secondly, in most sympatric sister species contrasts, LD among their ‘speciation loci’ was significantly higher compared to their respective genomic background LD (wilcox.test *p <* 0.001) (Fig. 4a). There was a discernible build-up of LD in older species pairs such as *Pundamilia* on Makobe and Ruti islands compared to younger species pairs of *Pundamilia* on Python and Kissenda islands (Fig. 4a), that emerged much more recently through hybrid parallel speciation (*77*).

**Fig. 4.**
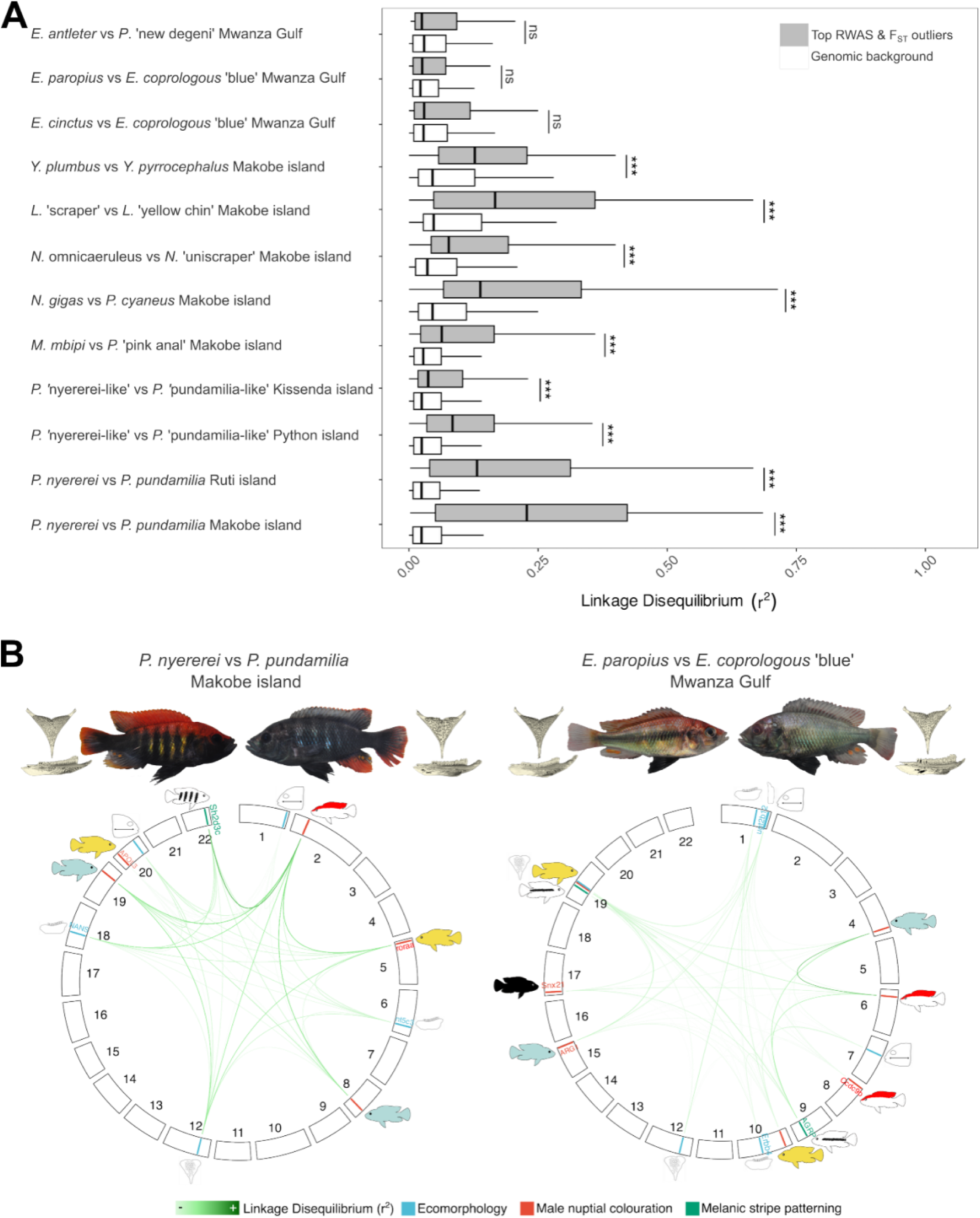
Coupling of ecological and mating trait modules during combinatorial speciation. (**a**) Linkage disequilibrium (LD; measured using *r*^2^ statistic) among RWAS top SNPs that are also top 5% F_ST_ (grey) outliers between closely related sympatric species compared to the genomic background LD (white). LD among top SNP is significantly higher than background LD comparisons (wilcox.test *p <* 0.001***) for all species pairs except demersal detritivores in the genus *Enterochromis*. (**b**) Genome-wide distribution of LD coupled alleles for two sympatric species pairs, one where LD among top SNPs is significantly elevated and one where it is not. It illustrates unique patterns of high interchromosomal LD among SNPs associated with ecological and mating traits that diverge in both of these speciation events but divergent in unique combinations among the species in each pair, consistent with expectation for combinatorial speciation from a redundant variant pool. The plots for the remaining species pairs are shown in Fig. S9. Lines on the ideogram represent a RWAS top SNPs that are also top 5% F_ST_ outliers; line colour denotes the trait complex the SNP was associated with in the RWAS: blue = ecomorphology, green = melanic stripe pattern, red = male nuptial colour motif as in Fig. 2; SNPs located in genes are annotated with gene names. Higher linkage disequilibrium (*r*^2^) values are depicted by darker green colours – see figure legend with scale. In all LD analyses only SNPs on different chromosomes and SNPs more than 100 Kbp away on the same chromosome were included to avoid effects of physical linkage. Even in the case of *Enterochromis* sister species where the LD among all top SNPs is not significantly elevated over the genomic background, the highest LD patterns among top SNPs map to trait combinations that differentiate these species.

Next, we compared the distribution of these ‘speciation loci’ across the genome and LD between them for each species-pair (Fig. 4b, Fig. S10). We find that loci contributing to ecological traits and mate choice traits are not physically linked in the genome but became coupled (presumably) during or shortly after speciation, with varying degrees of LD association among them. For example, between *P. nyererei* and *P. pundamilia* both trophic ecology (reef planktivore with fine pharyngeal bones and papilliform teeth vs crevice-dwelling insectivore with heavier pharyngeal bones and enlarged teeth) and male nuptial colour (flameback on yellow flank vs entirely blue flank) diverged, and this is reflected in the high LD between SNPs associated with these traits (Fig. 4b left). Similarly, *E. paropius* vs *E. coprologous* ‘blue’ are deeper water detritivores that diverged in male nuptial colouration (flameback on a light yellow flank vs blue flank with melanic underside), divergent melanic stripe patterns (present vs absent midlateral band), and subtle differences in trophic morphology (very fine pharyngeal bones and papilliform teeth vs slightly heavier pharyngeal bones and slightly enlarged teeth) which is reflected in LD among specific SNPs associated with these traits (Fig. 4b right). Long-range interchromosomal linkage disequilibrium between trait complexes in full sympatry suggests non-random mating among individuals with these divergent traits (i.e. speciation). Interestingly, we found that different and unique subsets of RWAS top SNPs had diverged in different species pairs, even among phenotypically similar species pairs (see *Pundamilia* species pairs from Makobe and Ruti versus Python and Kissenda, Fig. 4b, S7), reinforcing the idea that despite its extreme evolutionary youth, trait architectures can be characterised by redundant genotype-phenotype mapping in this radiation. The unique combination of LD patterns among subsets of trait-associated alleles in different species pairs that we observe provides support for the hypothesis of adaptive radiation through combinatorial speciation (*51*).

## Discussion

Paleontological research has shown that adaptive radiations have repeatedly punctuated the evolutionary history of biodiversity on Earth in bursts of diversification that defy expectations of gradual evolution(*78*). When adaptive radiation can proceed in the absence of geographical isolation among populations, it can in principle be a rapidly acting modulator of biodiversity in moments when ecological opportunity arises, i.e. when carrying capacity for species exceeds existing species diversity in a region (*79*). While this sometimes happens, it more often does not, and the conditions for it to happen defy current understanding. In a classical article back in 1954, Anderson & Stebbins proposed that such bursts of diversification can be enabled by hybridisation between species (*10*), a proposition that received support from botanists but long-standing criticism from zoologists (but see (*80*)). Yet zoologists have yet to resolve the puzzle of cichlid fish adaptive radiations in African Great Lakes: how did hundreds of ecologically and morphologically diverse species arise without geographic isolation and come to coexist in extraordinarily species rich assemblages within sometimes just a few thousand years? Here, for the first time we empirically demonstrate that key ecological and mating traits in this radiation correspond to independently inherited and often redundant genetic modules derived from hybrid ancestry, which are leveraged in combinatorial speciation (*51*). Combining modules for different traits in many different ways, like Lego bricks, or as Anderson & Stebbins (*10*) put it, elements from assembly lines of several car models co-produced in the same factory, made it possible for so many diverse species to evolve from a finite set of elements. We found that the large number of events of speciation in Lake Victoria in just a few thousand years are associated with the build-up and persistence in sympatry of long-range linkage disequilibrium among species-specific combinations of trait modules. Furthermore, we illustrate that genetic redundancy arising from parallel/convergent evolution of traits in the ancestors of the hybrid swarm and the subsequent combination of the parallel evolved architectures into a single hybrid population further increased flexibility for combinatorial speciation. We propose that modularity (independence) (*81*) in genomic architecture for different trait complexes that are key to ecological adaptation and mate choice was crucial for adaptive radiation to unfold at such a rapid pace through combinatorial speciation in the absence of geographical isolation.

Our association analysis revealed that all key traits associated with ecological diversification and speciation in the LV cichlid fish adaptive radiation are polygenic across the radiation and governed by a mix of some modest to larger effect alleles as well as many small effect alleles dispersed across the genome. However, while this pattern emerges across the radiation and is to a good extent due to genetic redundancy between species, divergence of the same traits in individual speciation events tends to be associated with only a subset of these genetic variants (Fig. 3b, Fig. S8, Fig. 4b, Fig. S9). This is also consistent with the observation that different trait-associated SNPs are recruited into high LD in different species pairs that diverge in the same set of traits (see flameback or pharyngeal keel depth in Fig 4b) and with the observation that the QTLs from laboratory crossing (*82*, *83*) are subsets of the variants associated with the corresponding traits in our radiation-wide association analysis. The compensatory phenotypic effects of genetically redundant alleles that we find add significantly to evolvability in key speciation traits in this adaptive radiation (*52*, *75*). The radiation-wide gene pool tends to hold multiple genetic solutions for species divergence in any particular combination of key traits through redundant genotype–phenotype mapping. Simulations have shown that polygenic and redundant trait architectures promote recurrent speciation and adaptive radiation in fusion-fission dynamics (*48*). Our analysis further revealed that the genetic architecture of speciation/species divergence is physically unlinked and non-pleiotropic i.e. it is modular (Fig. 4b, Fig. S9). It requires long range LD to build and then be maintained between multiple trait modules dispersed across the genome. This is consistent with earlier studies finding genome-wide divergence patterns already at very early stages of speciation in pairs of sympatric species (*22*, *84*). Taken together, we conclude that the global genetic architecture of key traits across the radiation is polygenic, but the genetic architecture of speciation is oligogenic. It is conceivable that moderate selection would be sufficient to maintain LD underlying oligogenic architectures during non-geographic speciation. In contrast, polygenic architectures would require strong selection on speciation traits, which we do not see evidence for in the field as strong selection would constrain both speciation and trait diversity and speciation. Redundant genetic architectures of speciation may also allow emerging species to retain some genetic polymorphisms even for those traits that fixed under selection during speciation. Such context-dependent conversion of adaptive variation into cryptic variation and the reverse may facilitate successive speciation events in the same lineage involving divergence in (some of) the same trait sets.

The low pleiotropy and absence of physical linkage across adaptive radiation trait architectures may be surprising at first. Pleiotropic or clustered (i.e. physically linked by inversion polymorphisms) architectures were often suggested to facilitate speciation with gene flow. However, such architectures would constrain diversification beyond a finite number of ecotypes or species, as has been observed in sympatric and parapatric ecotypes of stickleback, and to some extent in Darwin’s finches (*85–87*). The reason for this is that clustered architectures provide low evolutionary modularity – few components that can evolve independently without resulting in correlated effects on linked traits (*88*), and thus strong genetic constraints to the dimensionality of phenotypic evolution and low evolutionary flexibility. For the phenotypic complexity required for coexistence of large number of species through resource partitioning and mate choice to evolve rapidly, it seems advantageous if genetic variants affect only one trait at a time and are independently inherited (*89*). The modular nature of both mating and ecological traits that we demonstrated here for Lake Victoria cichlids may therefore be key to promoting diversification into many phenotypically diverse and unique species within the same and in different environments (*51*). This is consistent with classical theory on the importance of evolutionary modularity in promoting diversity in natural systems by circumnavigating constraints of selection on other traits (*90*). However, much of the empirical work on modularity is limited to morphological modularity and its role in diversification and adaptive radiation (*91–94*). Our study illustrates that both modularity in traits (illustrated by low phenotypic covariance among traits across the radiation (Fig 2a) and many different combinations of essentially the same set of craniofacial traits, dentition traits, body stripe patterns, and male nuptial colour motifs) and in their underlying genomic architecture is necessary for rapid diversification of a lineage into many sympatric species. The low pleiotropy among trait alleles that is required for this, may be explained by the enrichment of ancient admixture variation, originally derived from distantly related species that hybridized and then maintained through several fusion and fission cycles in the history leading to this radiation. Low pleiotropy would be unexpected had many of these trait-associated alleles had emerged from *de-novo* mutation within the radiation.

While we focused on SNPs here, it is very likely that small inversions or other small structural variants contribute additionally to modular trait architectures (*70*). In the cichlid radiation in Lake Malawi, large inversions that were exchanged between distantly related clades by hybridisation have been found to facilitate parallel adaptation to the same environmental gradients between these clades (*95*). While this explains parallel ecological transitions between major habitats it cannot explain how hundreds of ecologically diverse species arose within each major habitat. As we do not uncover any evidence for physical linkage among speciation and adaptive radiation trait modules in Lake Victoria cichlids, inversion polymorphisms that almost certainly segregate also in this radiation (*95*, *96*) do not appear to have captured major genetic variants that would physically couple key mate choice and adaptation traits in the radiation.

Questions that remain in order to understand the process of explosive adaptive radiation in Lake Victoria concern the nature of the evolutionary forces that generate and maintain various patterns of long-range linkage disequilibrium between genetic modules in the absence of geographical isolation. As important as the absence of strong genetic correlations between adaptive radiation traits is, the absence of strong ecological correlations between these traits, which we can also infer from our trait correlation matrix (Fig. 2A), is equally important. We argue this implies the absence of consistent selection constraint on trait associations. We show that trait values at ecological and mating traits can be combined in many ways and that some alleles associated with these two categories of traits in any specific sympatric species pair tend to be in elevated long-range LD. We interpret this as a signal of non-random mating among sympatric species, something for which good behavioural evidence also exists (*58*, *62*). In the presence of gene flow, the strength of phenotypic divergent selection required on a trait or trait combination to maintain LD increases with the number of loci involved in phenotypic divergence which can be at least partly offset by non-random mating (*97*). Oligogenic speciation architectures as observed here might require only very moderate strengths of selection in the face of non-random mating. Clearly, LD among multiple traits, many involving more than one region in the genome, including traits that are associated with mate choice, persists in sympatry without physical linkage. Future work will have to address how it arises from a situation of no LD (or very different LD) in the first place. One possibility is initial build-up of a novel pattern of LD among ecological and mating traits happens in transient geographical or ecological parapatry, and that the extent of assortative mating generated in this way becomes quickly sufficient for incipient species to persist as they reinvade each other’s ranges and form sympatric species communities. Our findings reconcile quantitative genetics of traits with population genomics of speciation and have broad implications for understanding conditions under which adaptive radiation can act as a modulator of biodiversity when ecological opportunity arises. With the genomic architecture of species divergence in ecological and mating traits for many species revealed, future work can address the remaining questions, such as the evolutionary forces driving LD patterns, through combining ecological and genomics work at the scale of entire assemblages of species.

## Acknowledgements

Many thanks to the Tanzania Fisheries Research Institute (TAFIRI) for hosting OS and his team during fieldwork from 1989 to now. We are very grateful for the technical support of M. Haluna and M. Kayeba during the fieldwork of OS and his team in Mwanza over the last three decades since 1989. We thank COSTECH for research permits and the Tanzania Department of Fisheries for export permits. We thank the Research group of OS for discussion and feedback. Many thanks also to the members of the Seehausen laboratory and C. Bank laboratory at the University of Bern for the many excellent discussions. We thank A. Nolte for the DNA extraction protocol, J. Johnson for fish illustrations, and M. Koller for figure editing support. We thank P. Nicholson and the NGS platform of the University of Bern, Switzerland, for sequencing services and support, and we thank N. Zemp and the entire GDC at ETH Zürich, Switzerland, for HPC support. We remember our colleague David Alexander Marques who was a PhD student and Postdoc of OS before he became curator of birds at the Natural history of Basel. He studied the role of hybridisation in radiations of fish and birds. David sadly passed away on 7 January 2026 just weeks before this paper was submitted.

## Author contribution

Pooja Singh: conception, lead all analyses and interpretation, and manuscript writing. Heidi Tschanz-Lischer: genome assembly and annotation. Kassandra Ford: 3D morphometrics data collection and analyses. Ehsan P. Ahi: candidate gene & GO analysis interpretation. Marcel Haesler: fish husbandry and tissue dissections.

Salome Mwaiko: DNA and RNA extraction. Joana I. Meier: DNA extraction. David Marques: conception. Remy Bruggman: genome assembly support. Mary Kishe: fieldwork. Ole Seehausen: fish collection and identification, conception, interpretation and manuscript writing.

## Data, code, and materials availability

Pundamilia nyererei v3 genome assembly and annotation and the raw PacBio Hi-C and PacBio Iso-seq data is deposited to the NCBI SRA (BioProject PRJNA1231206). Individual resequencing data for 313 individuals is sourced from (*9*, *70*, *98*), with the exception of 11 newly sequencing sampled that have been deposited to NCBI SRA BioProject PRJNA1231206. All sample and phenotyping information is in File S1. 3D scans have been deposited to BioImageArchive (doi: 10.6019/S-BIAD2928). Scripts used in this study are available on Github: https://github.com/poojasingh09/2025_singh_radiation_gwas.

## Supplementary Materials

Materials and Methods Supplementary Text

Table S1

Figures S1 – S11

File S1

