## Supplemental Text, Figures, Table for "Modular and redundant genomic architecture underlies combinatorial mechanism of speciation and adaptive radiation"

Science
AAAS

### Supplementary Materials for

**Modular and redundant genomic architecture underlies combinatorial mechanism of speciation and adaptive radiation**

Pooja Singh *et al*

**The PDF file includes:**

Supplementary Results
Tables S1
Figs. S1 to S11
Materials and Methods
References

**Results**

**Candidate genes controlling ecological and mating traits are dispersed across the genome**

For all 14 traits we identified many genic as well as non-coding SNPs dispersed widely across the genome, including in well-known genes and previously unknown candidate genes (Fig. 3a shows results for five traits, see Fig. S3 for all 14 traits). Most strikingly, both **midlateral** and **dorsolateral band** shared an intergenic top SNP on chromosome 5 that was highly associated with their presence (MLB $\beta$=0.15, PIP = 0.74 and DLB $\beta$ = 0.13, PIP = 0.99). Dorsolateral band never occurs in this radiation without the midlateral band, and this SNP may regulate this interaction. The top genic SNP for midlateral band with an effect size of $\beta$ = 0.12 (PIP > 0.48) was in the *agrp* (*asip2*) gene that is known through QTL analysis and Crispr-cas9 as a genetic switch for midlateral band formation in cichlids(*1*), as mentioned above. A top genic SNPs for dorsallateral band was in *cyp27a1*, known as a vision gene(*2*) and the top SNP for **vertical bars** mapped to *sez6* that is interestingly associated with exploratory behaviour and cognition in mice(*3*).

The four highest top SNPs for flameback include two **flameback** associated QTLs previously found in a cross of *P. nyererei*-like (a flameback species) and *P. pundamilia*-like (no flameback), with the region on chromosome 2 being a QTL for red dorsum and the region on chromosome 8 for red dorsal fin(*4*). Candidate genes for the flameback male nuptial colour were vision: *slc5a8*(*5*) and carotenoid/melanin/iridophore associated: *rarb*(*6*), *ahr*(*7*), *alkal2*(*8*). The top flameback gene *plxdc2* has not been associated with pigmentation previously but plexins are implicated in melanocyte regulation(*9*). Gene Ontology (GO) analysis of flameback top SNPs found enrichment ($q < 0.05$) of 'retinoic acid receptor signalling pathway' that regulates chromatophores(*10*), 'neurotransmitter secretion' that controls physiological colour change in vertebrates(*11*) and 'regulation of Toll signalling pathway' that is implicated in melanogenesis(*12*) (Fig. S4). **Yellow flank** and **blue flank** are two reciprocally exclusive male nuptial colour motifs known to be repeatedly involved in speciation (*13*). An intergenic top SNP on chromosome 2 for blue flank overlaps with a QTL for yellow flank in a cross of *P. nyererei*-like (has yellow flank) and *P. pundamilia*-like (has blue flank)(*4*). The strongest candidate gene for both these traits was *pdgfrb*, a gene thought to be involved in fish colouration as it evolved from a duplication of the proto-*kit/csf1r* gene when fish acquired chromatophores(*14*). This may suggest that yellow and blue flank can sometimes be alternative allelic states of the same gene. Other skin/vision pigment-related candidates for yellow flank are *sox5* which is a molecular switch for cell fate in xanthophores (*15*), *roraa*(*16*), *rarb*(*6*), and *clul1*(*17*). Candidates for blue flank coloration are *clul1*(*17*) and *tead1*(*18*), the latter of which regulates *mitf*, a master regulator of melanocyte differentiation and animal pigmentation(*19*). Blue flank top SNPs were also enriched for 'hippo signalling' (Fig. S4) that is involved in retinal differentiation(*20*) and melanogenesis(*21*). Interestingly, a top genic SNP with positive effect on male **melanic body** colouration mapped to *nos1ap*, a gene associated with social status switches in cichlids(*22*). Melanic body had melanogenesis associated candidates *trpv4*(*23*), *wnt10b*(*24*), *adamts*(*25*); and more broadly melanic body top SNPs were enriched for the 'Wnt signalling pathway' that is involved in the differentiation and proliferation of melanocytes, as well as the regulation of melanin production(*26*). Top candidates for **red chest** male colouration were *smyd3*, a methyltransferase that regulates mesoderm development where chromatophores are situated(*27*, *28*). Overall, gene associations for male nuptial colour motifs were either involved in pigmentation or vision. The latter observation supports the idea that sexual selection by female mate choice and male-male competition contribute to speciation and species differentiation in Lake Victoria cichlids(*29–31*). Male nuptial colouration can be associated with water depth and ambient light conditions in cases of speciation(*32*, *33*) and the evolution of female preferences for male nuptial colouration likely involves interactions between sexual selection(*34*), ecological adaptation in the visual system to water depth/light(*33*), frequency dependent selection by male-male competition(*35*) and possibly other components of behaviour that remain to be identified.

We found many significant SNPs/genes associated with craniofacial shape **PC1** and a few for PC2 (Fig. 3). The strongest predictor of a species' position on craniofacial shape PC1 (the bite force versus suction feeding continuum, with short vs longer heads and jaws) is a top SNP on chromosome 21 located near the *vsx* (alias *rinx1*) gene that is involved in craniofacial anomalies and modulating interpupillary distance in humans (*36*). Interestingly, *lrriq* gene that is physically linked to *alx1* is an important locus for beak shape in Darwin's finches and great tits (*37*, *38*) and also one of the top candidate genes for craniofacial PC1 in our analysis (chromosome 2). Another candidate gene for PC1, *Efemp1* controls head length in zebrafish(*39*). Other candidates known to be involved in cichlid bone/cartilage/muscle development were *fgf14*, *mybph*, *col8a2*(*40*) and the sex-specific development gene *foxo3*(*41*). The intergenic top SNP for **PC2** on chromosome 20 overlaps with a QTL for preorbital depth in a cross between *Neochromis omnicaeruleus* and *P. nyererei*-like (*42*). A promising candidate for shape variation along **PC2** (body depth, cheek depth and eye size continuum) is *roraa* that is involved in orofacial cleft phenotype in humans(*43*). The top candidate genes for **cusp shape** were *macf1*, which controls tooth mineralisation in mice(*44*); *lrp6* that interacts with the wnt pathway during tooth development in humans(*45*) and *prdm16* that is required for normal tooth development in mice(*46*). *Ugt1a* was the strongest candidate for **number of inner tooth rows** in the upper oral jaw. This gene has not previously been associated with dentition. Another candidate gene, *fbp2*, overlaps with a QTL on chromosome 22 for this trait in a cross between *N. omnicaeruleus* (3 to 8 inner rows of teeth) and *P. nyererei*-like (2 inner rows of teeth) (*42*). Lower pharyngeal jaw **keel depth** top SNPs mapped to many bone development genes such as *hk2*, *mapk14a*, *wnt7a* and *nans*. The latter is required for human bone development in several craniofacial regions, including the pharyngeal arch skeleton (*47*). The top candidate gene for **pharyngeal tooth molarisation** was *vmp1*, which has not been previously associated with tooth development. However, another candidate for this trait was *pknox2*, a transcription factor that plays a crucial role in the early stages of tooth development in mammals (*48*).

**Table S1 Comparison of long-read improved assembly and annotation of Pundamilia nyererei (v3.0) in this study compared to previous versions.**

| | Brawand et al 2014 (v1.0) | Feulner et al 2018 (v2.0) | Singh et al (v3.0) |
|---|---|---|---|
| **Quality** | Unanchored, short-read assembly & RNAseq annotation | v1.0 anchored with linkage map | Anchored, PacBio improved genome & Iso-seq annotation |
| **Nb scaffolds** | 7,236 | 6,876 | 2,201 |
| **N50** | 2.5 Mb | 29.8 Mb | 31.3 Mb |
| **Total length** | 830.1 Mb | 856.2 Mb | 916.1 Mb |
| **Totel length (without N)** | 698.8 Mb | 698.8 Mb | 890.9 Mb |
| **Genes** | 24,222 | 24,222 | 29,621 |
| **Isoforms** | 40,489 | 40,489 | 54,586 (with functional annotation) |

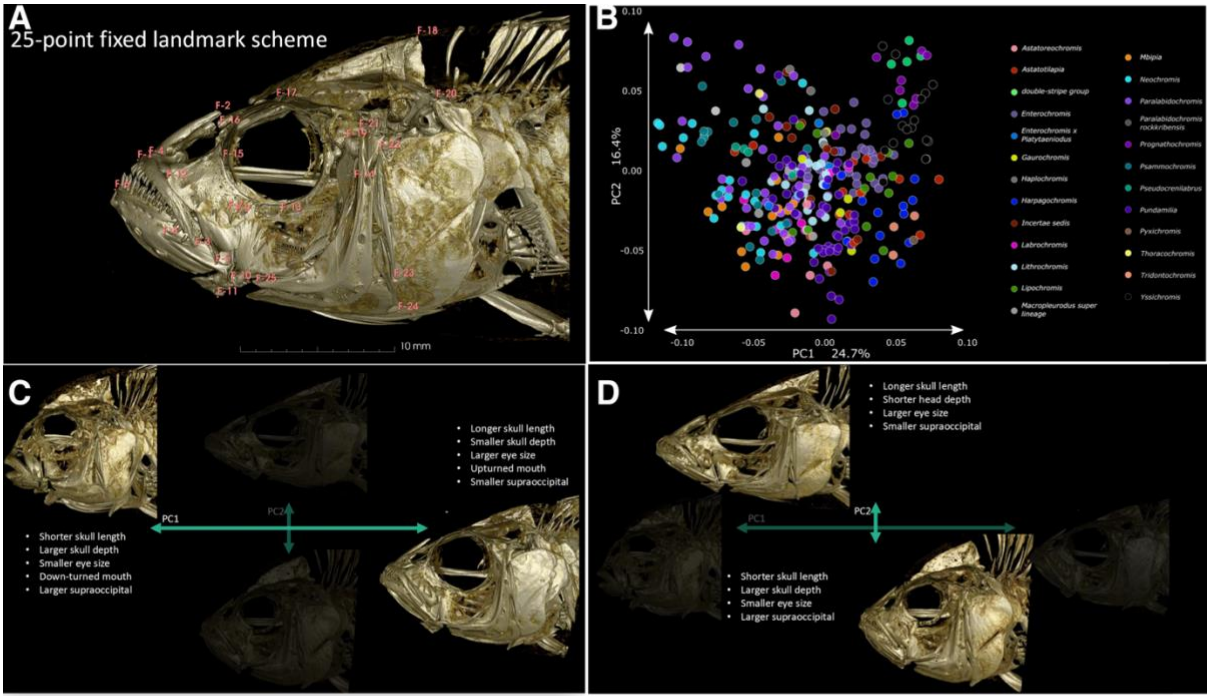

**Fig. S1 Overview of 3D geometric morphometric analysis of craniofacial shape.**
(a) Principal component analysis of 3D craniofacial geometric morphometric analysis shows major variation in head shape. (b,c) Traits contributing to variation in Principal components 1 and 2.

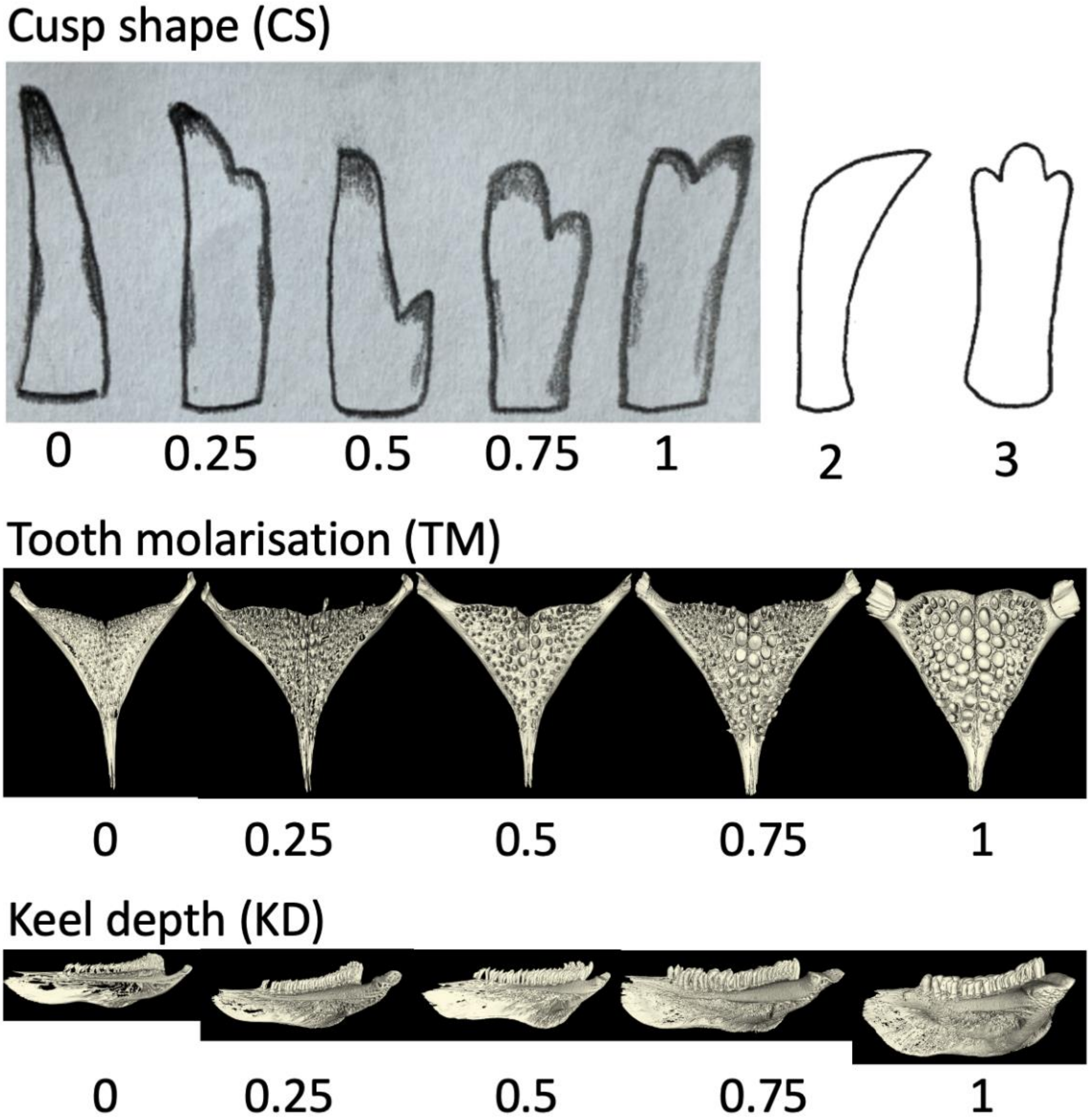

**Fig. S2 Trait categories for craniofacial ecomorphology traits oral teeth cusp shape, lower pharyngeal jaw tooth molarisation, and lower pharyngeal jaw keel depth.**

Number of inner rows of upper oral teeth not shown as the categories were the number of rows.

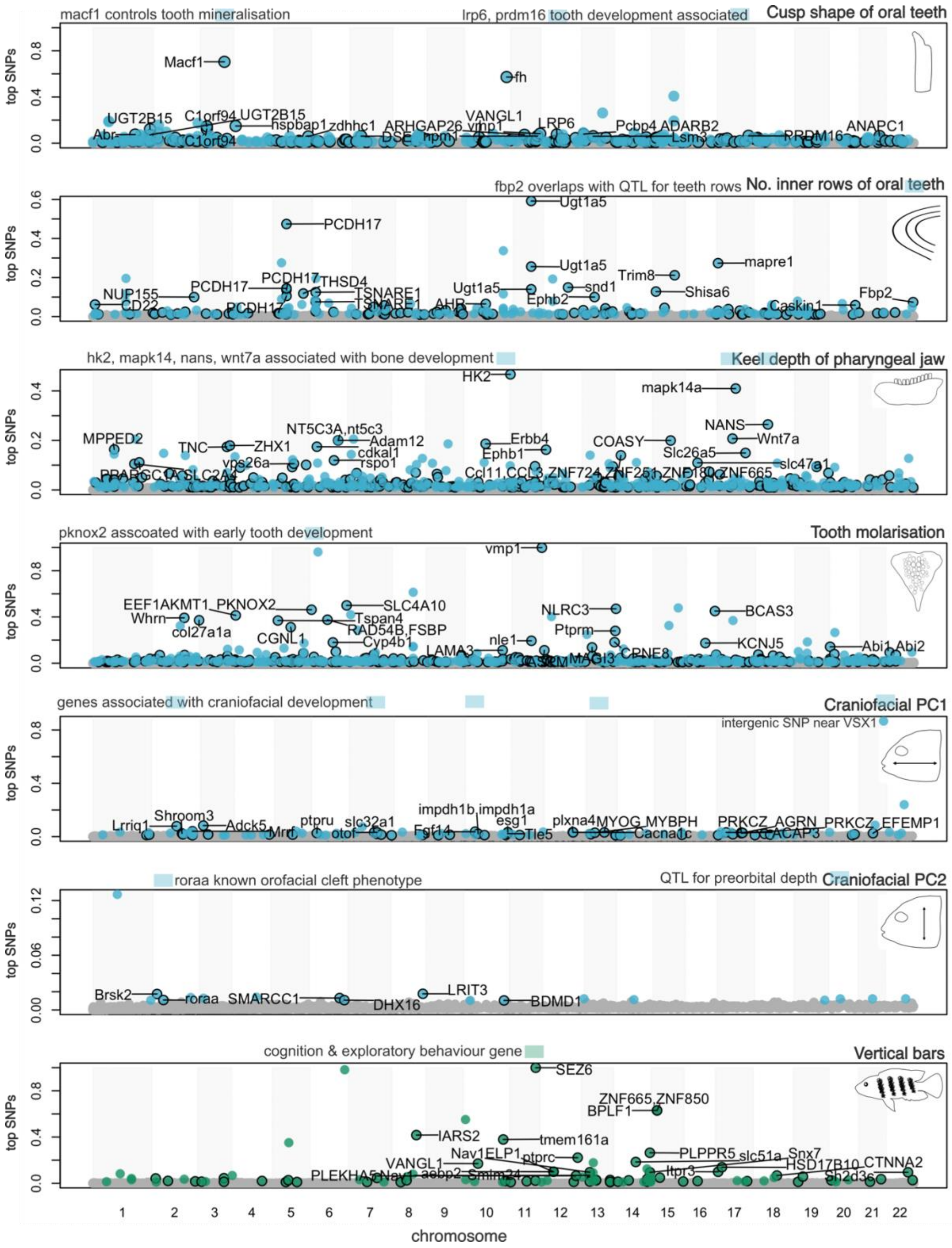
macf1 controls tooth mineralisation
lrp6, prdm16 tooth development associated
Cusp shape of oral teeth
top SNPs
Macf1
fh
UGT2B15
C1orf94
UGT2B15
hspbap1
zdhhc1
ARHGAP26
vmp1
VANGL1
LRP6
Pcbp4
ADARB2
Abr
C1orf94
DSE
hpm1
Lsm3
PRDM16
ANAPC1
fbp2 overlaps with QTL for teeth rows
No. inner rows of oral teeth
Ugt1a5
PCDH17
Ugt1a5
mapre1
Trim8
snd1
Shisa6
Fbp2
NUP155
CD22
PCDH17
PCDH17
THSD4
TSNARE1
TSNARE1
Ugt1a5
AHR
Ephb2
Caskin1
hk2, mapk14, nans, wnt7a associated with bone development
Keel depth of pharyngeal jaw
HK2
mapk14a
NANS
Wnt7a
MPPED2
NT5C3A,nt5c3
Adam12
cdkal1
rspo1
TNC
ZHX1
vps26a
Erbb4
Ephb1
COASY
Slc26a5
slc47a1
Ccl11,CCL3,ZNF724,ZNF25,ZNF180,ZNF665
pknox2 asscoated with early tooth development
Tooth molarisation
vmp1
EEF1AKMT1
PKNOX2
SLC4A10
Tspan4
RAD54B,FSBP
Whrn
col27a1a
CGNL1
Cyp4b1
NLRC3
Ptprm
BCAS3
KCNJ5
Abi1,Abi2
LAMA3
nle1
MAGI3
CPNE8
genes associated with craniofacial development
Craniofacial PC1
intergenic SNP near VSX1
Lrriq1
Shroom3
Adck5
ptpru
slc32a1
otof
impdh1b,impdh1a
esg1
Fgf14
Tle5
plxna4
MYOG
MYBPH
Cacna1c
PRKCZ
AGRN
PRKCZ
ACAP3
EFEMP1
roraa known orofacial cleft phenotype
QTL for preorbital depth
Craniofacial PC2
Brsk2
roraa
SMARCC1
DHX16
LRIT3
BDMD1
cognition & exploratory behaviour gene
Vertical bars
SEZ6
ZNF665,ZNF850
BPLF1
IARS2
tmem161a
Nav1
ELP1
ptprc
VANGL1
PLEKHA5
Nav1
asbp2
Smim24
PLPPR5
slc51a
Snx7
Itpr3
HSD17B10
CTNNA2
Sh2d3c
chromosome
1 2 3 4 5 6 7 8 9 10 11 12 13 14 15 16 17 18 19 20 21 22

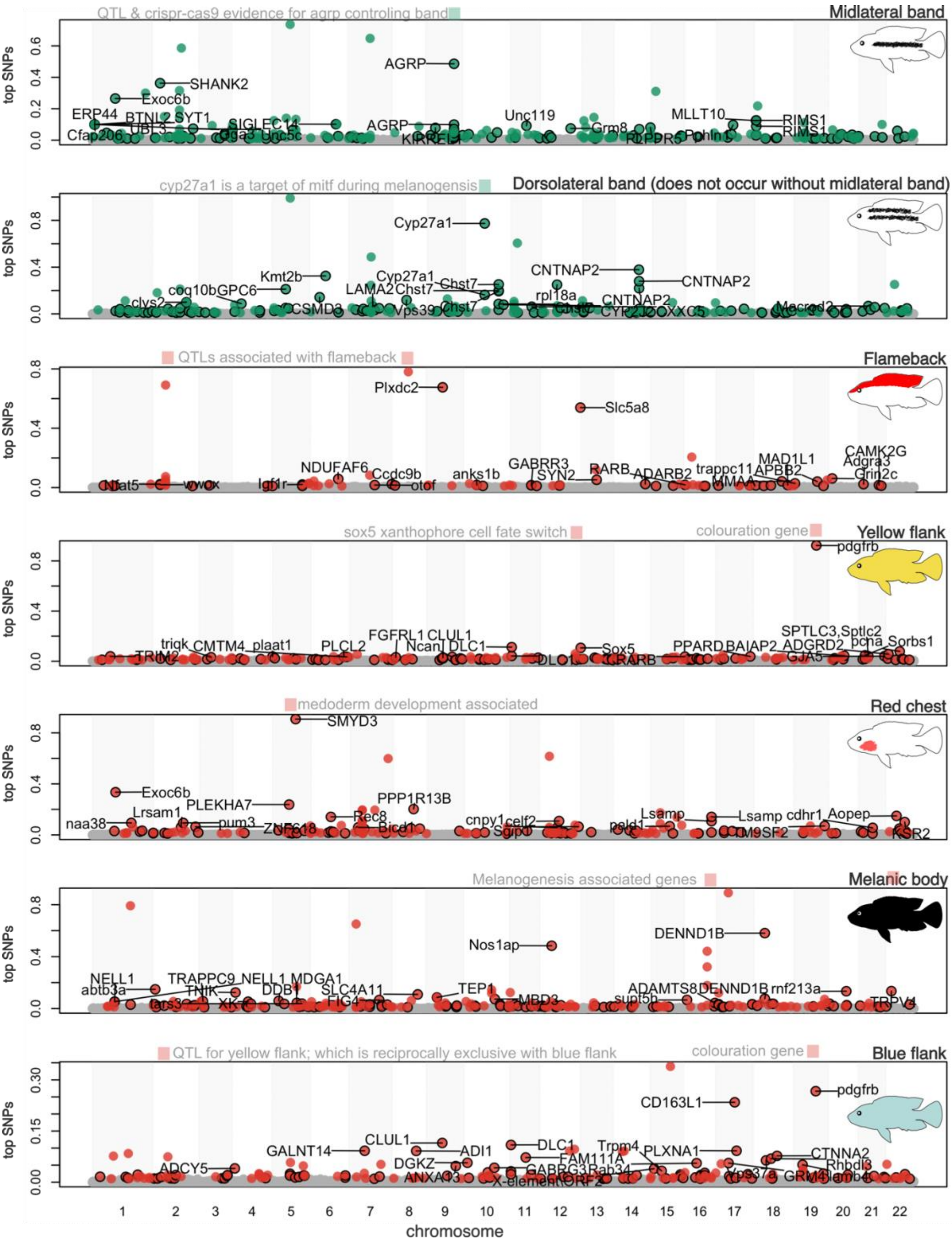

**Fig. S3 Top RWAS SNPs for 14 traits are dispersed across the genome.**

Significant top SNPs are highlight as coloured circles and non-significant SNPs are denoted by grey circles. Colour of circles are categorised by the trait complex that the trait belongs to: blue-ecomorphology, green-melanic stripe patterning and red-male nuptial colour. Coloured circles with black outline are SNPs located in genes and coloured circles without black outline are SNPs located in intergenic regions. Top 20 genic SNPs are annotated with gene names.

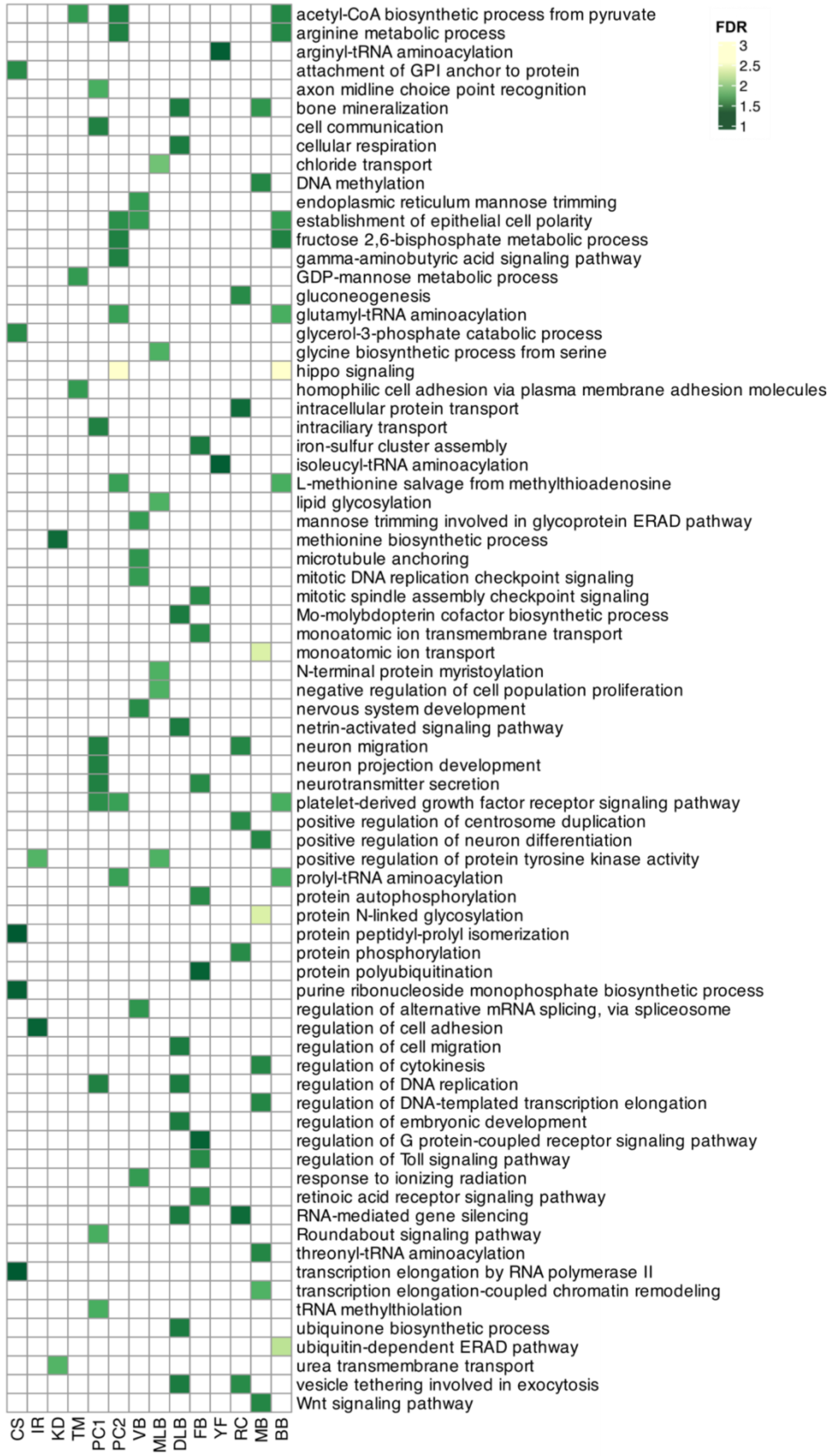

**Fig. S4 GO enrichment (only FDR < 0.05 shown) of top RWAS SNPs.**

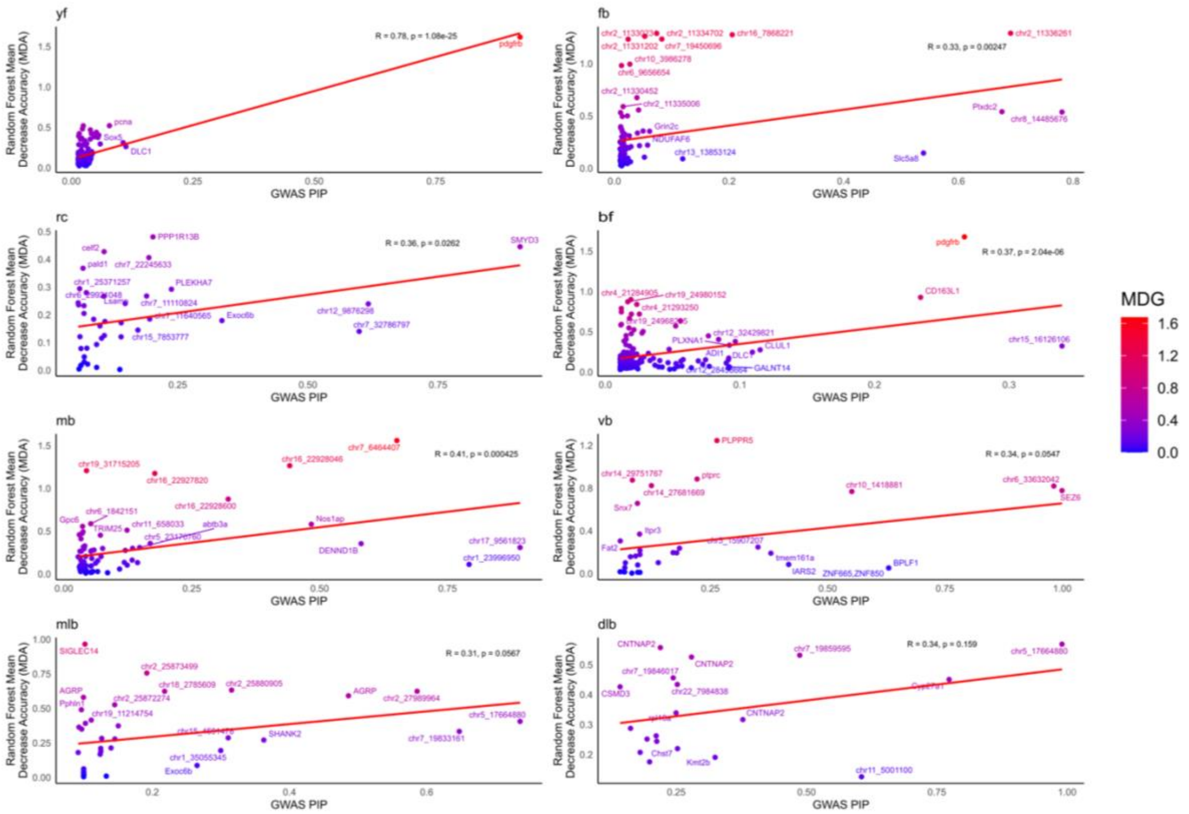

**Fig. S5 Machine learning predictive accuracy of top SNPs for binary body colour and patterning traits using random forest models.**

For most traits, a significant Pearson correlation exists between RWAS top SNP PIP score and Mean Decrease in Accuracy (MDA) from random forest decisions trees. MDA is a Random Forest variable importance measure that tells you how important a predictor (SNP) is for prediction accuracy. Higher MDA means the SNP was important for accurately predicting the presence/absence of the trait. Mean Decrease in Gini (MDG), a secondary Random Forest measure is illustrated for each SNP as a colour heatmap. MDG measures how much a SNP reduces Gini impurity on average, across all trees. Higher MDG means the SNP is more frequently and more effectively used to split the data.

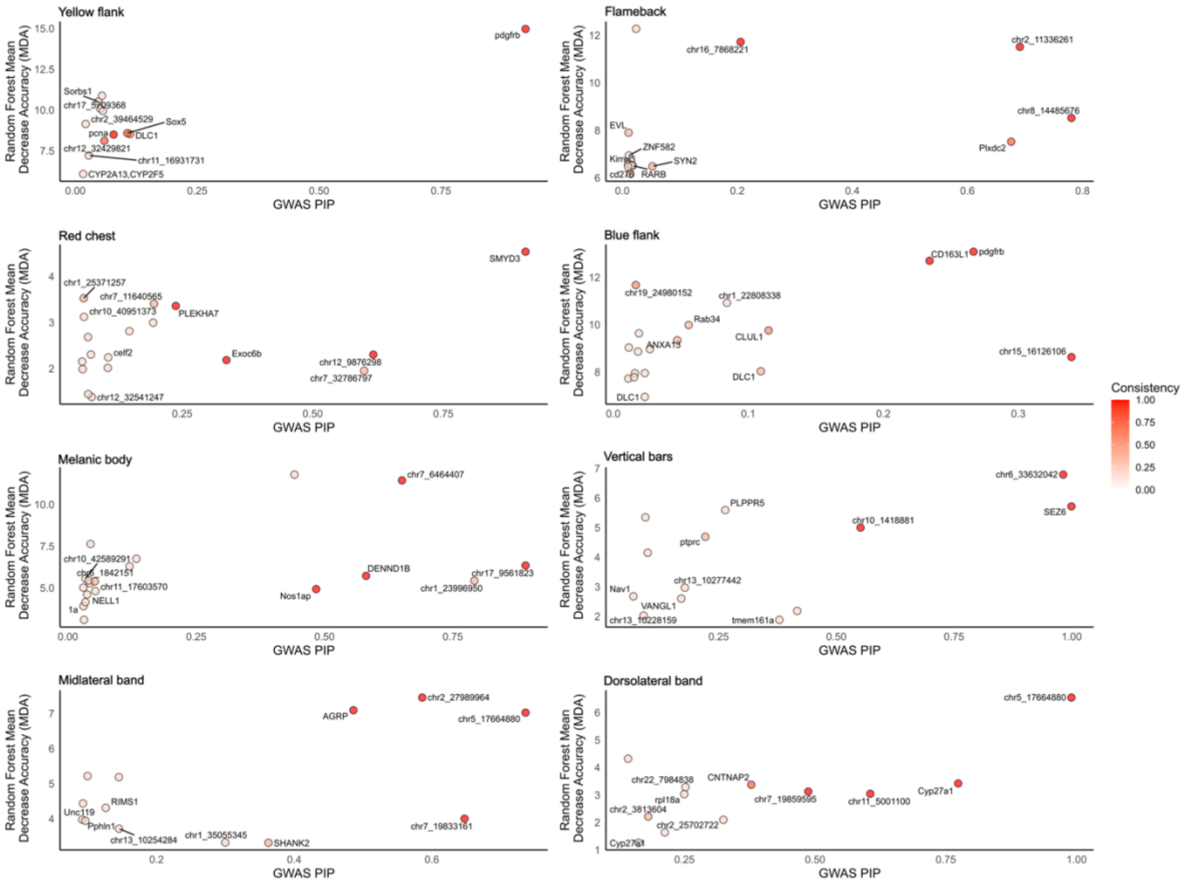

**Fig. S6 Consistency of machine learning predictive accuracy of top SNPs for binary body colour and patterning traits using random forest models.**

Top SNPs with high consistency were robust genomic predictors for the trait across . Consistency = Number of Folds where SNP was Selected / Total Number of Folds.

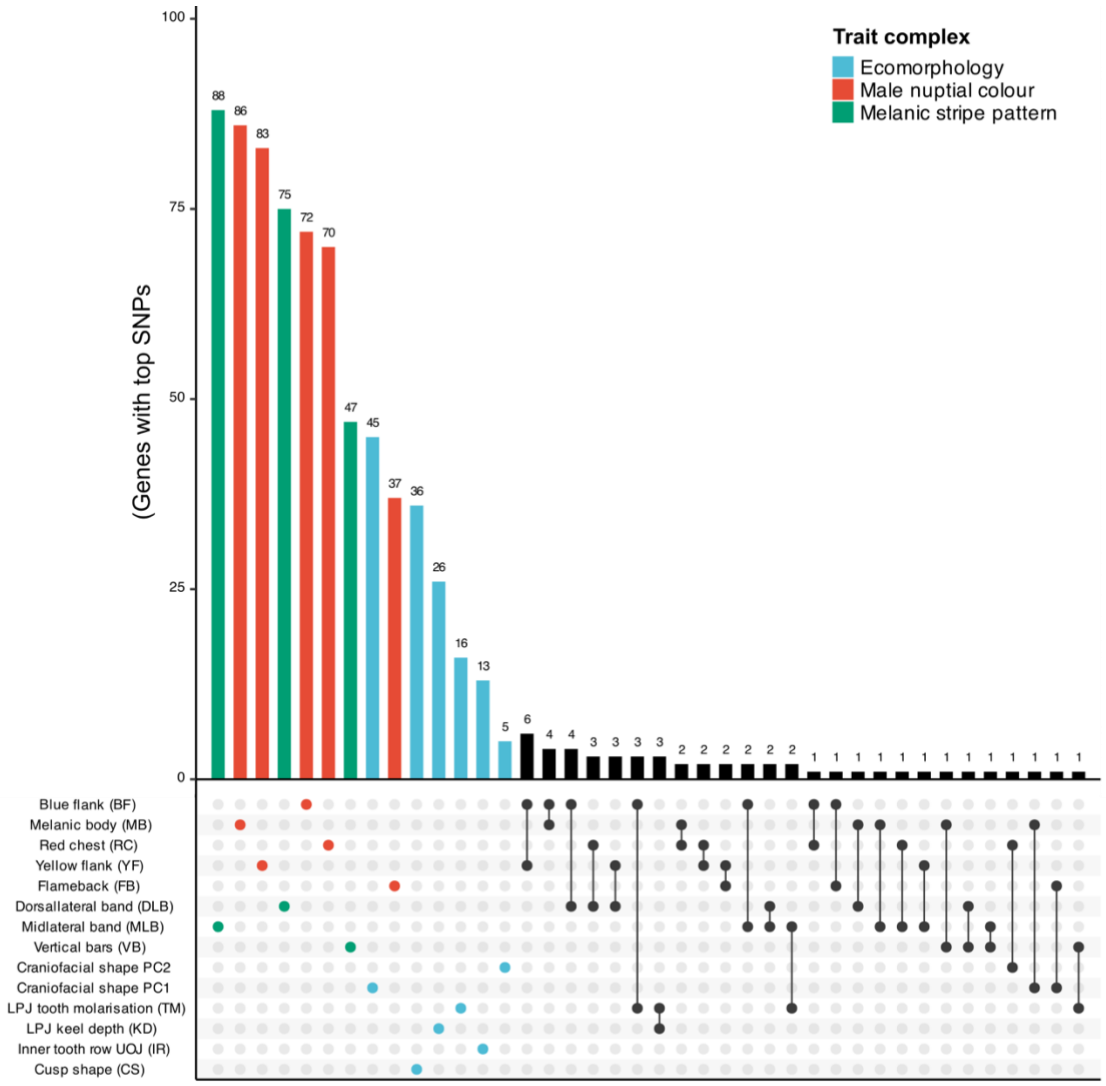

**Fig. S7 Overview of genes the top SNPs from the RWAS map to. Number of overlapping genes across traits are shown. Trait bars are coloured by the trait complex that they belong to.**

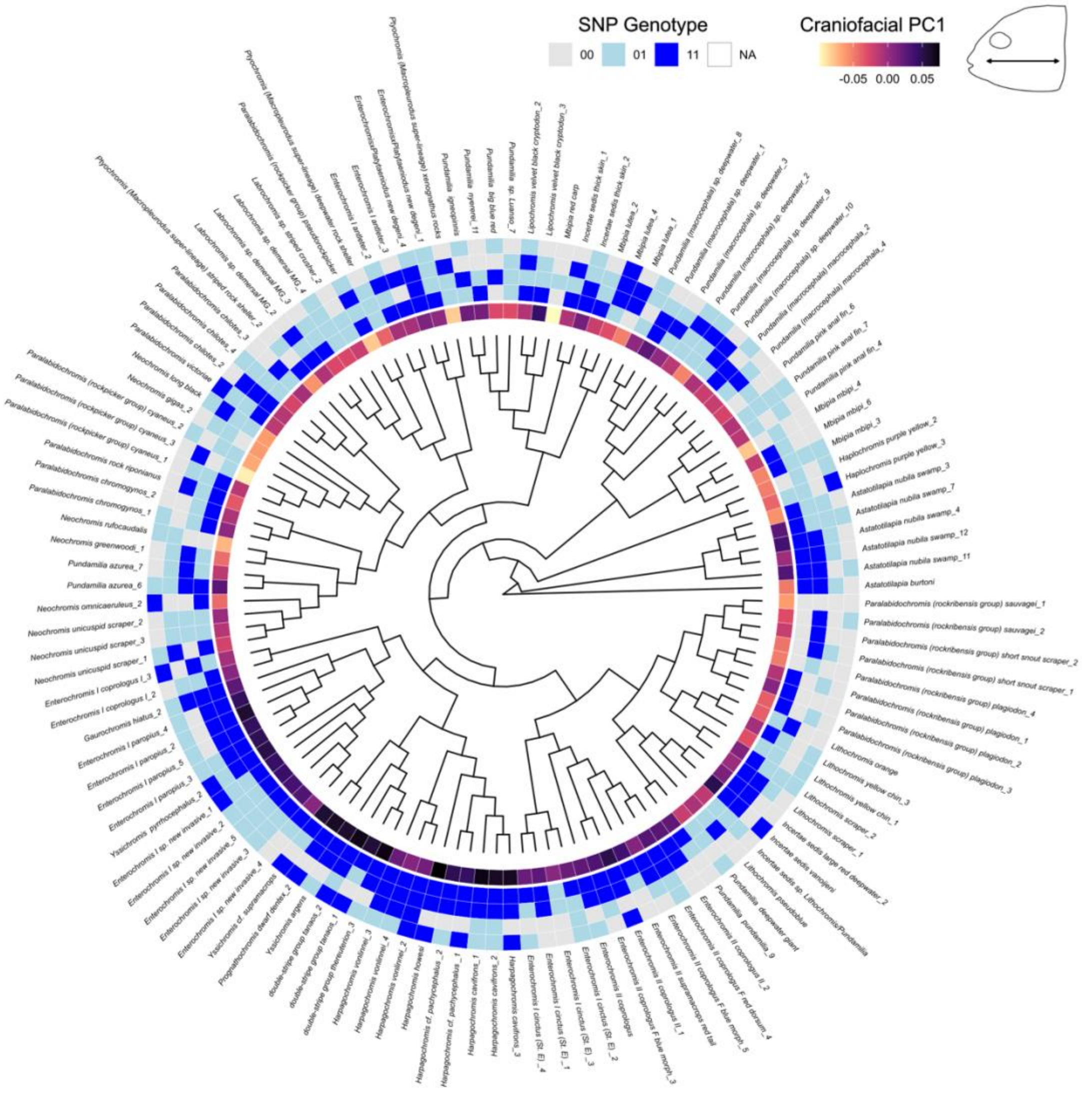
SNP Genotype
00
01
11
NA
Craniofacial PC1
-0.05 0.00 0.05

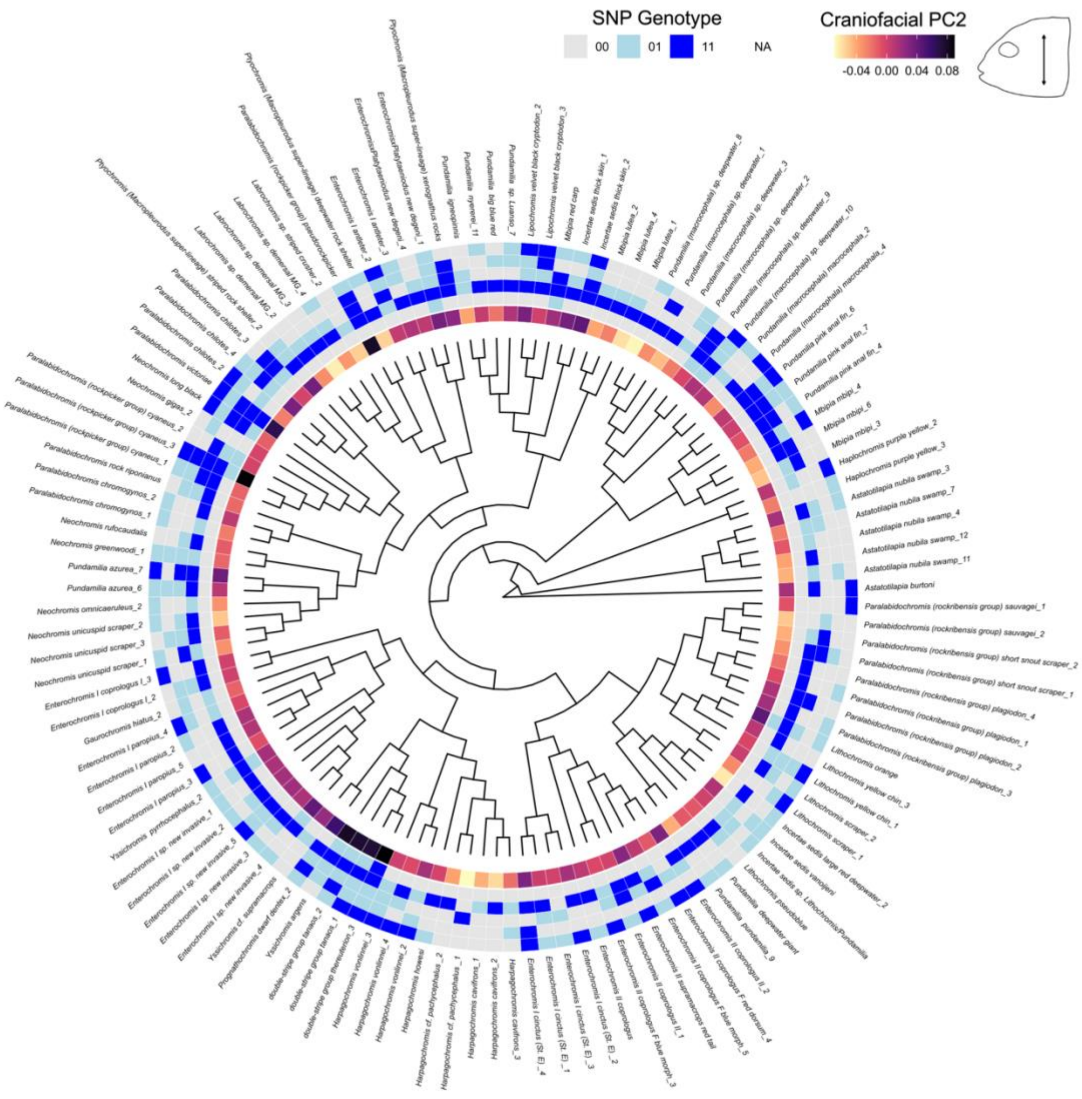
SNP Genotype
00
01
11
NA
Craniofacial PC2
-0.04 0.00 0.04 0.08

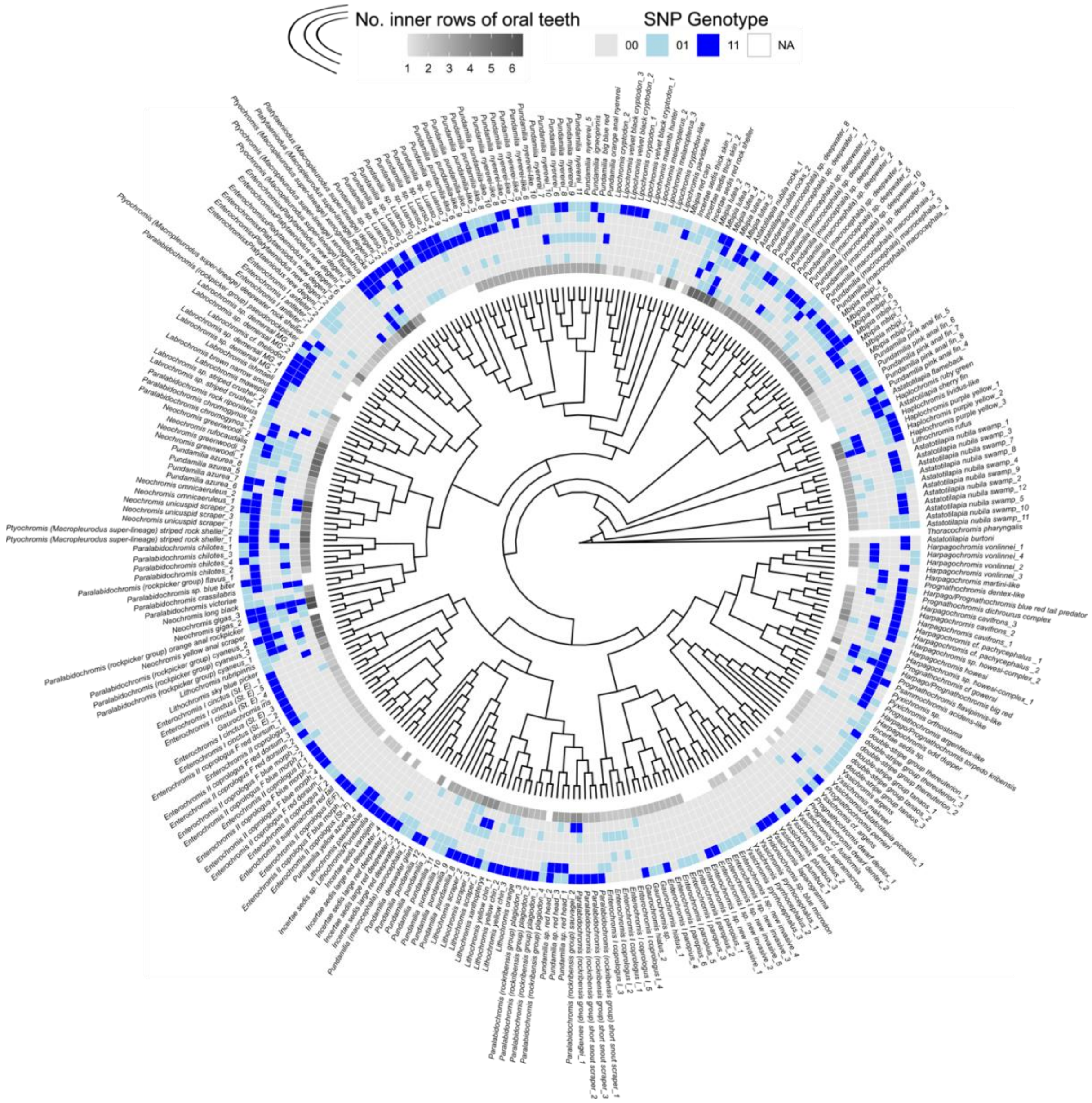
No. inner rows of oral teeth
1 2 3 4 5 6
SNP Genotype
00 01 11 NA

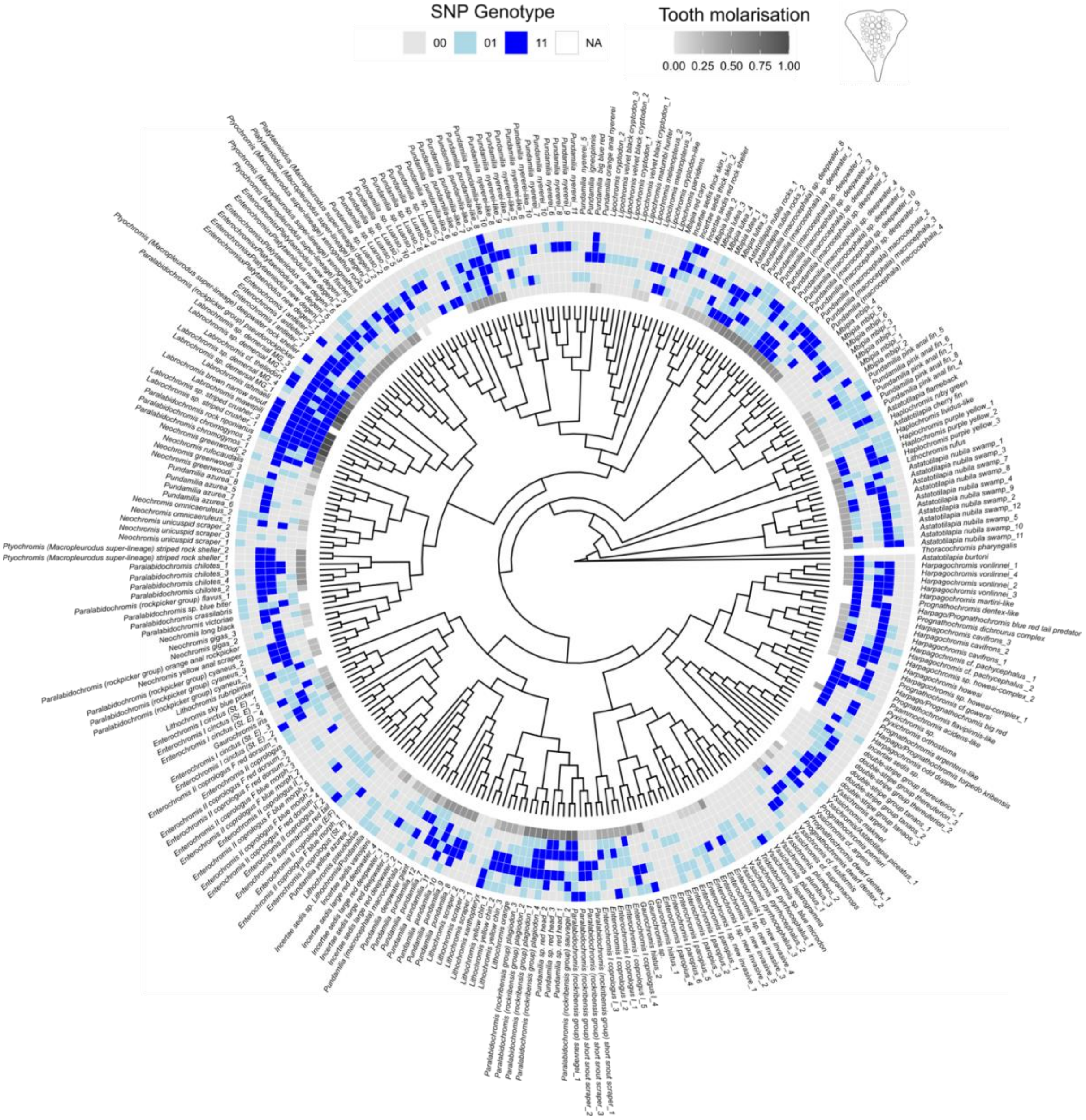
SNP Genotype
00
01
11
NA
Tooth molarisation
0.00 0.25 0.50 0.75 1.00

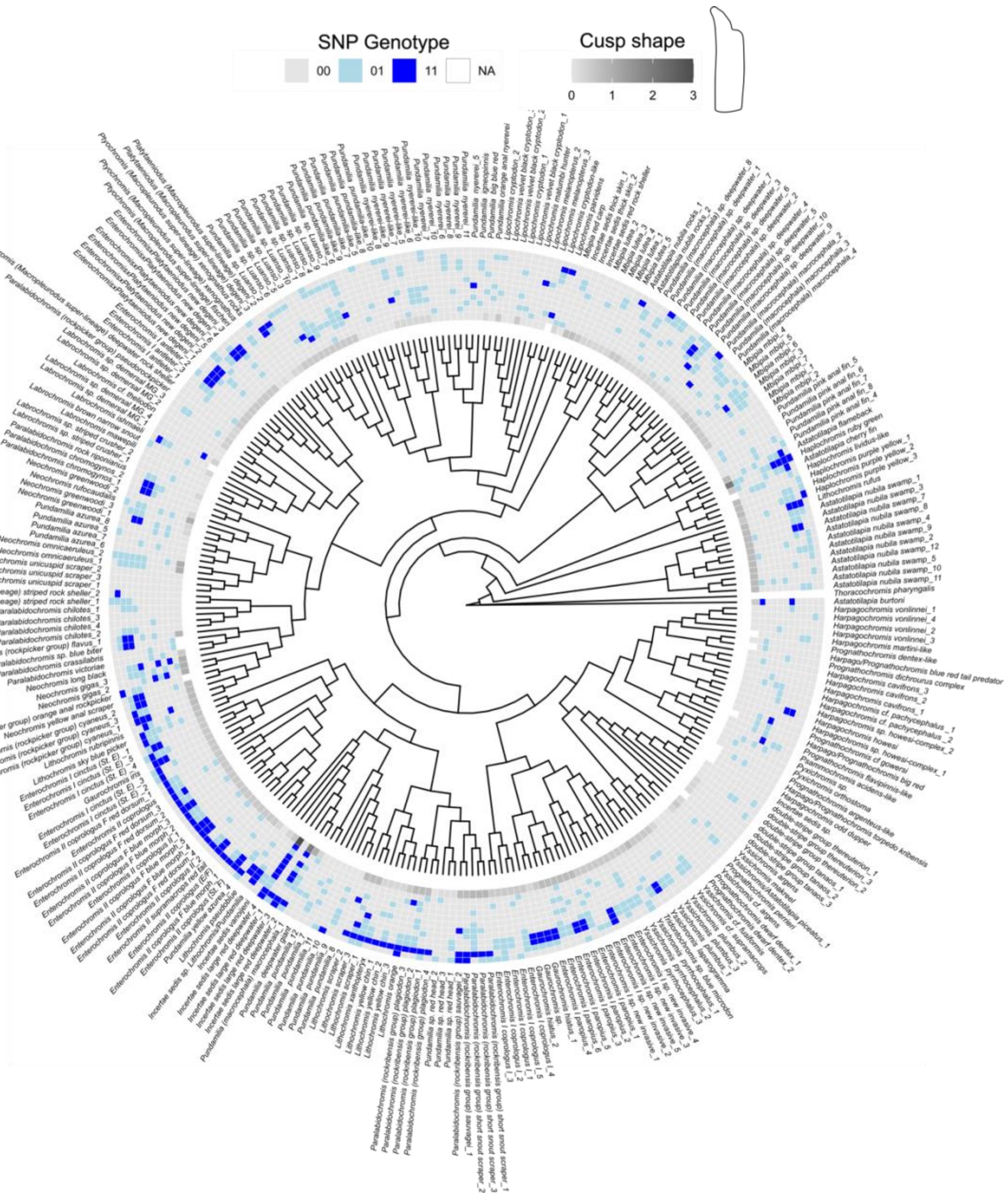
SNP Genotype
00
01
11
NA
Cusp shape
0
1
2
3

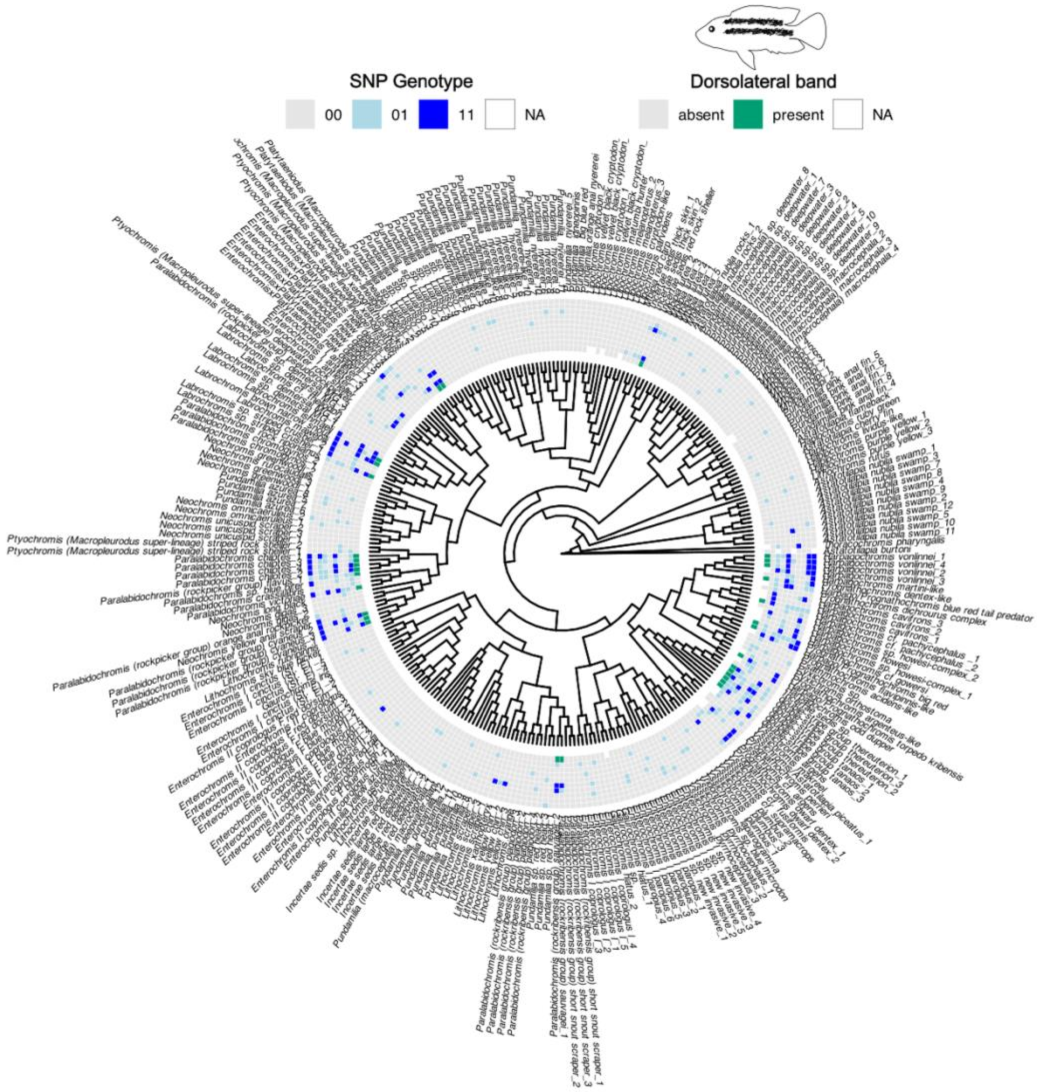
SNP Genotype
00
01
11
NA
Dorsolateral band
absent
present
NA

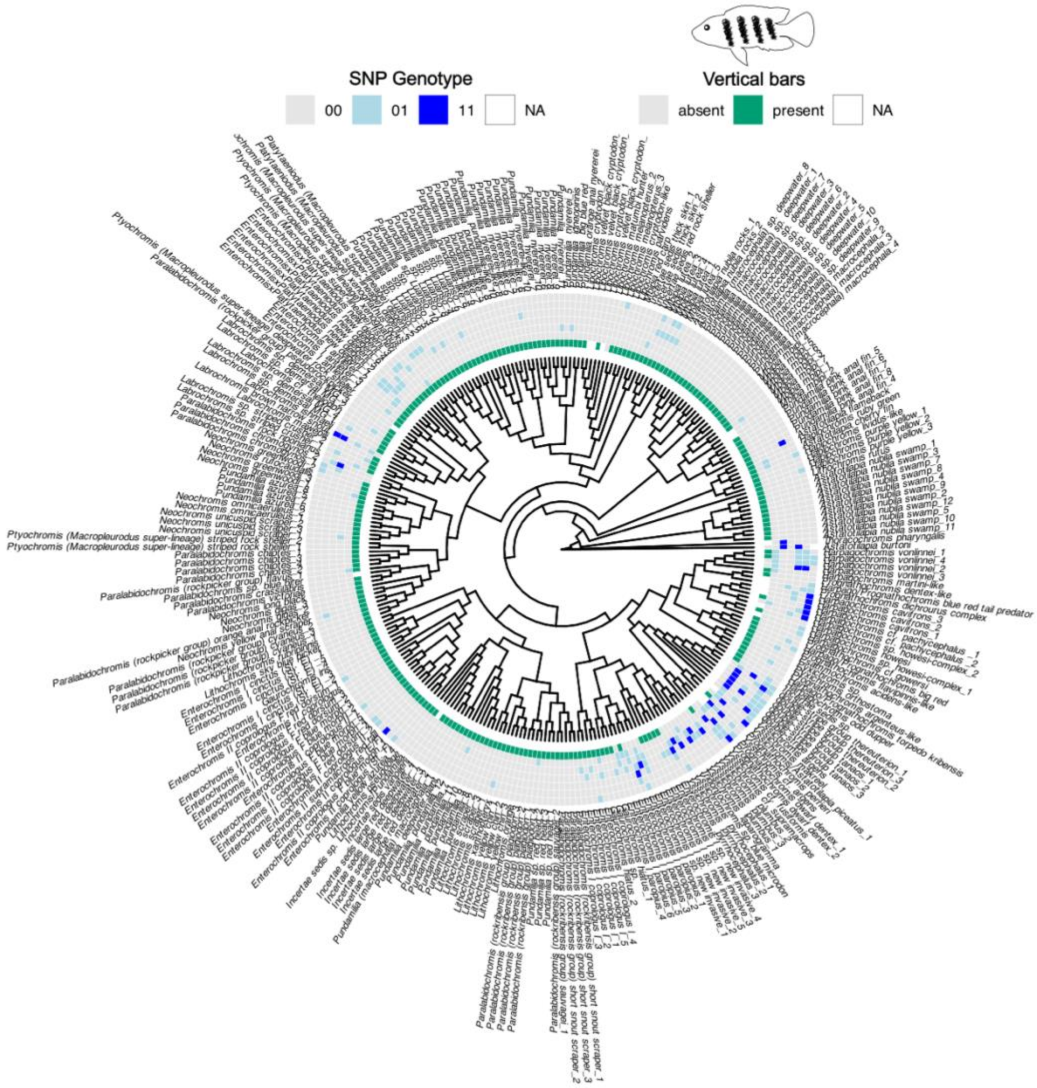
SNP Genotype
00
01
11
NA
Vertical bars
absent
present
NA

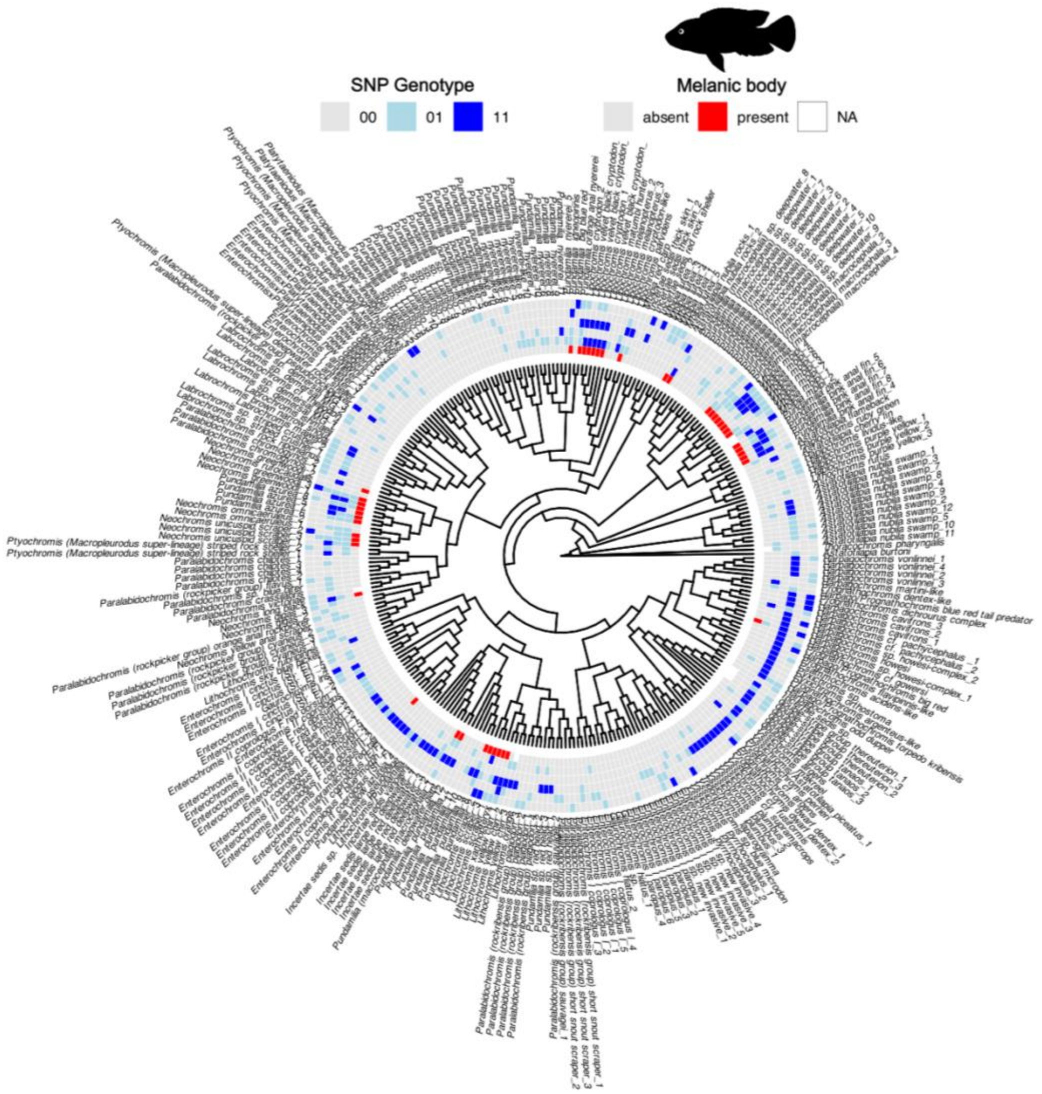
SNP Genotype
00
01
11
Melanic body
absent
present
NA

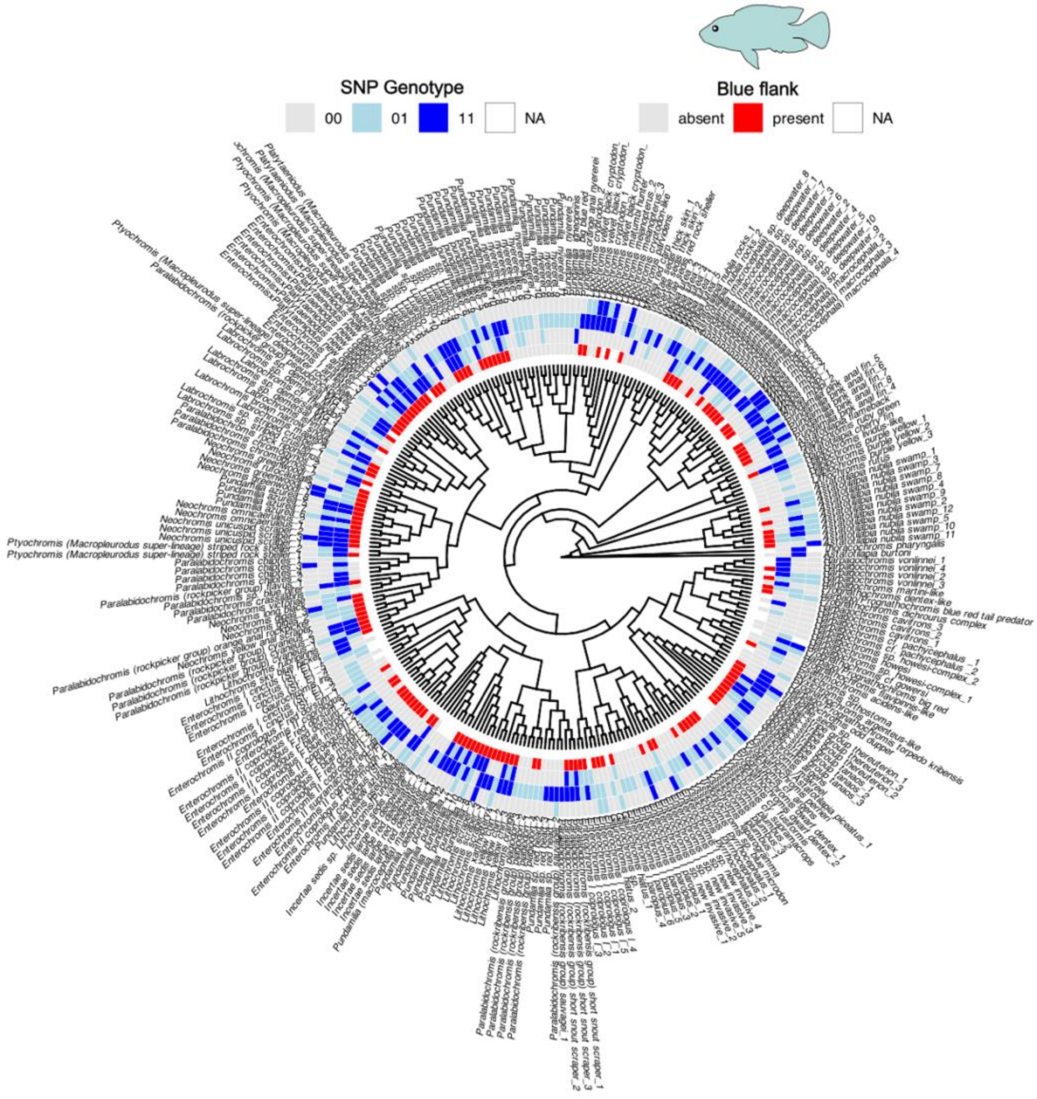
SNP Genotype
00
01
11
NA
Blue flank
absent
present
NA

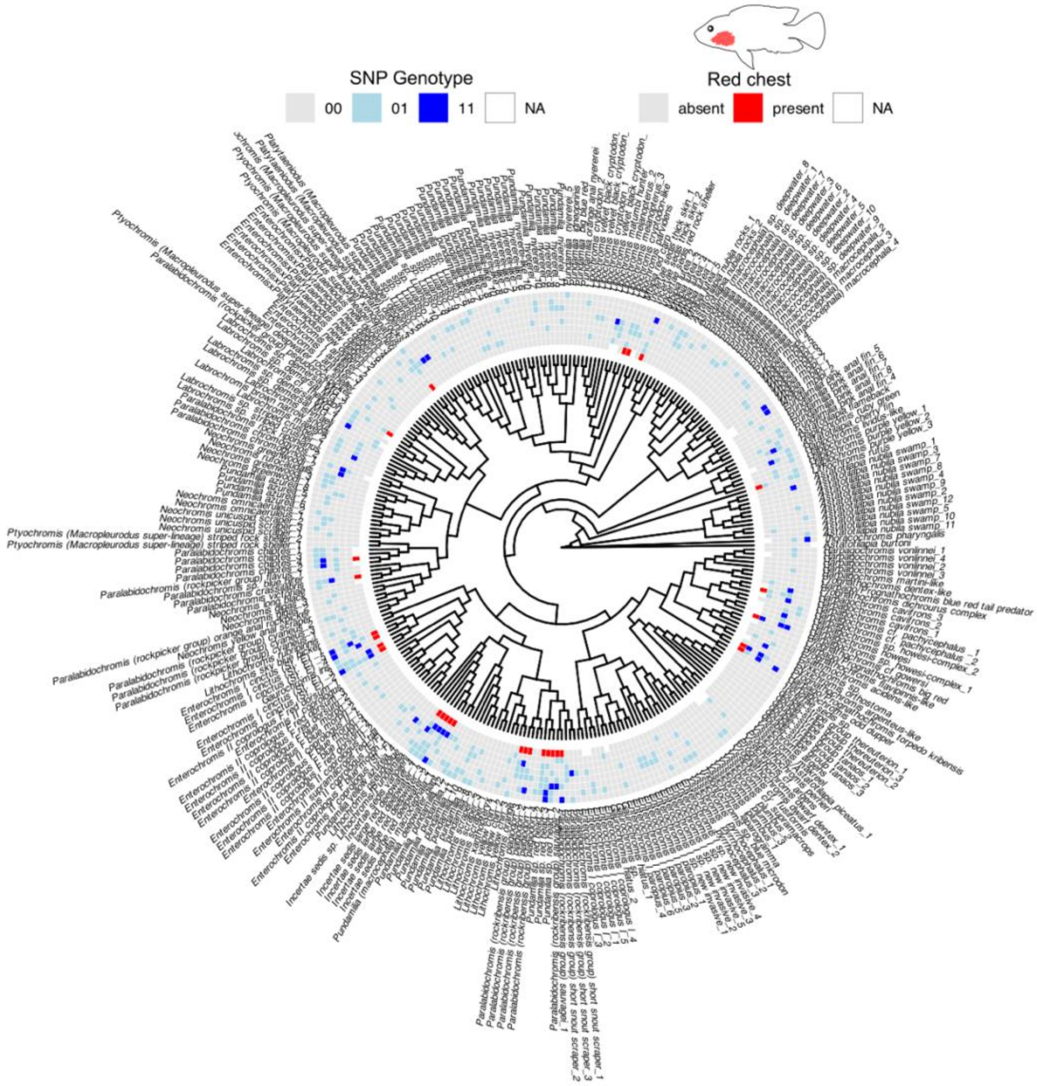
SNP Genotype
00
01
11
NA
Red chest
absent
present
NA

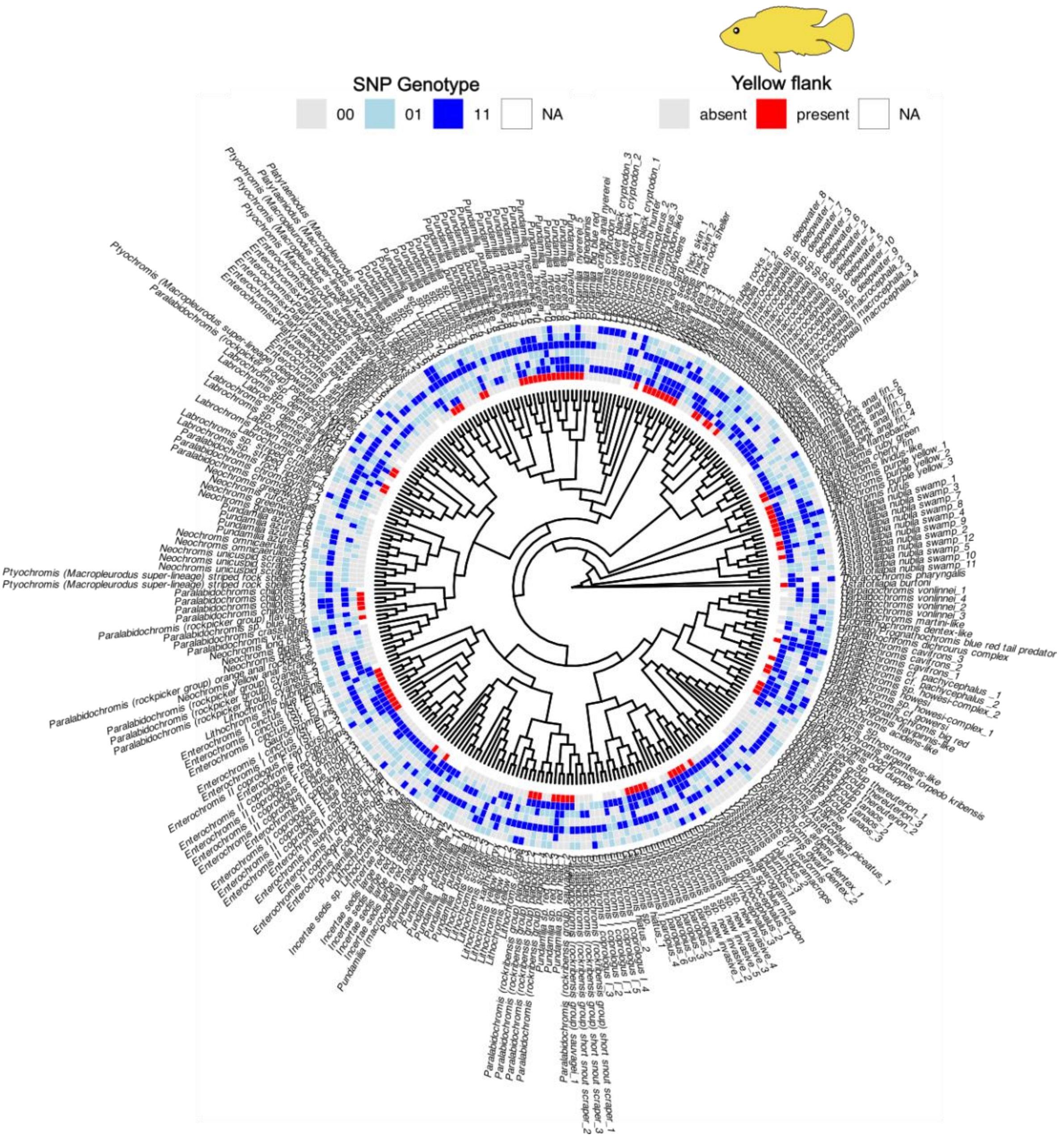
SNP Genotype
00
01
11
NA
Yellow flank
absent
present
NA

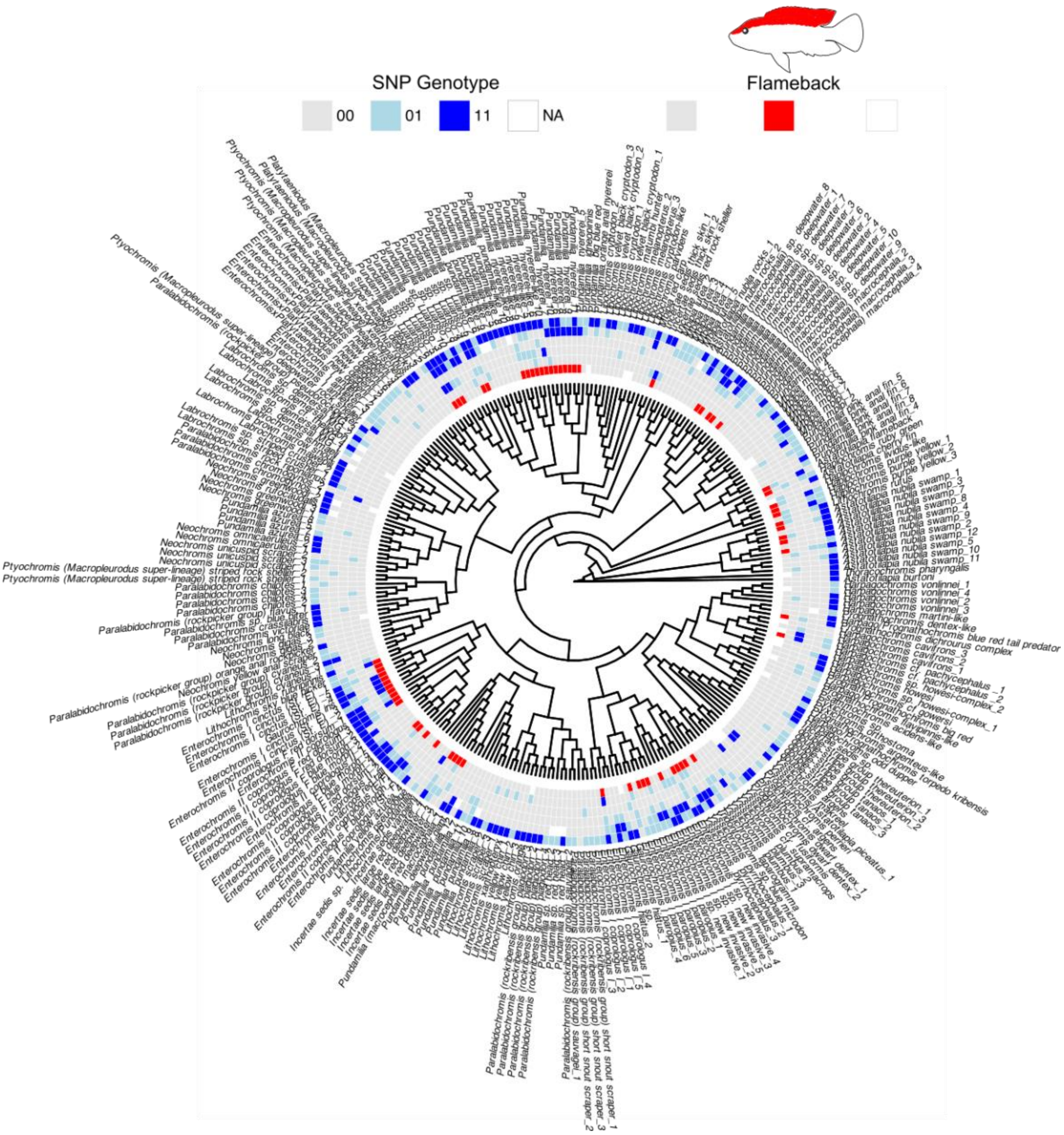

**Fig. S8 Genetic redundancy illustrated across the radiation for all traits except Midlateral band and Keel Depth which can be found in main Fig. 3.**

Genotypes of select top SNPs from the RWAS are illustrated for all individuals in this study along with the phenotype. Genome-wide SNP based phylogeny from Fig. 1 is used.

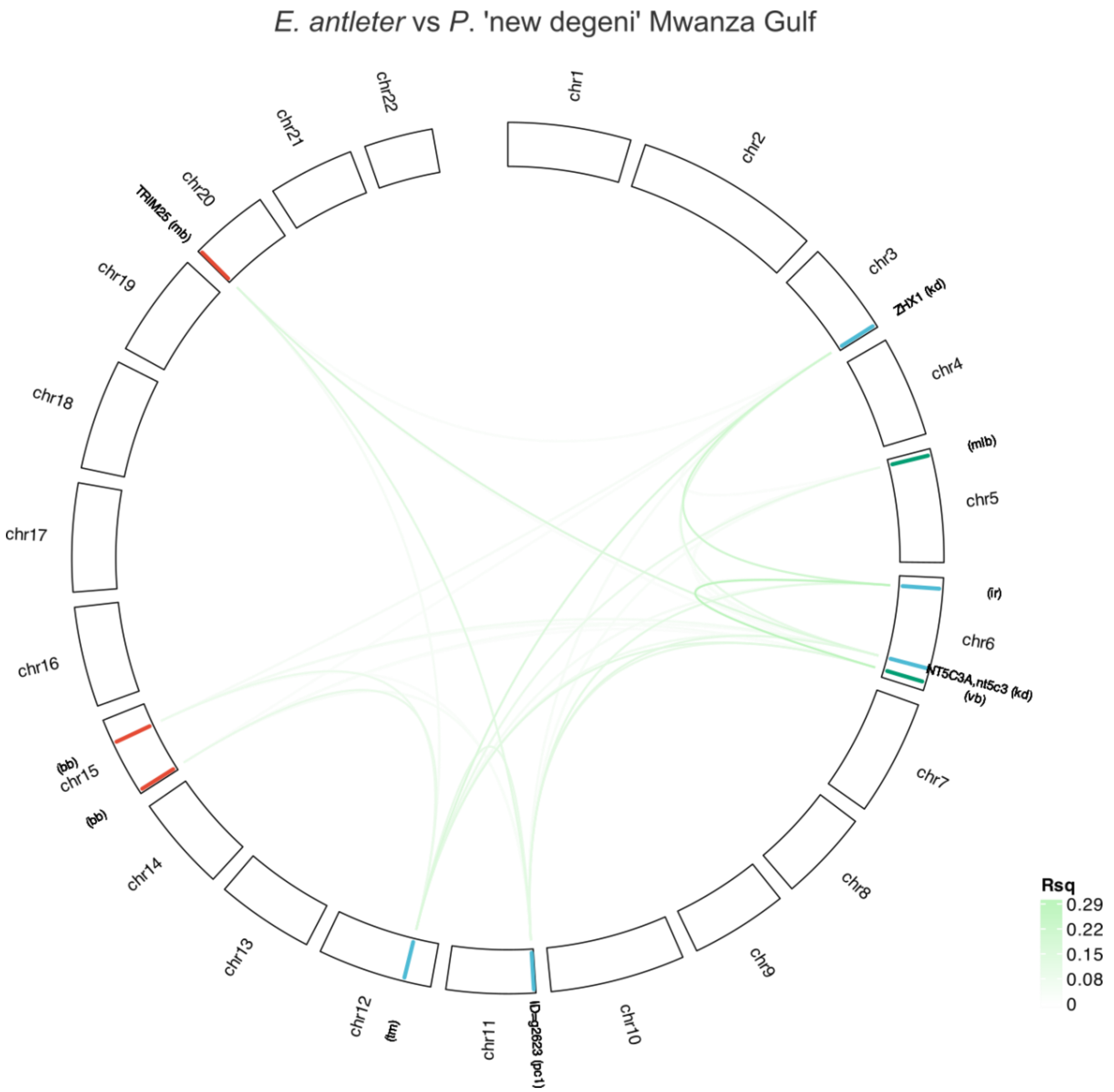
*E. antleter* vs *P.* 'new degeni' Mwanza Gulf
chr1
chr2
chr3
ZHX1 (kd)
chr4
(mlb)
chr5
(ir)
chr6
NT5C3A,nt5c3 (kd)
(vb)
chr7
chr8
chr9
chr10
ID=g2623 (pc1)
chr11
(tm)
chr12
chr13
chr14
(bb)
chr15
(bb)
chr16
chr17
chr18
chr19
TRIM25 (mb)
chr20
chr21
chr22
Rsq
0.29
0.22
0.15
0.08
0

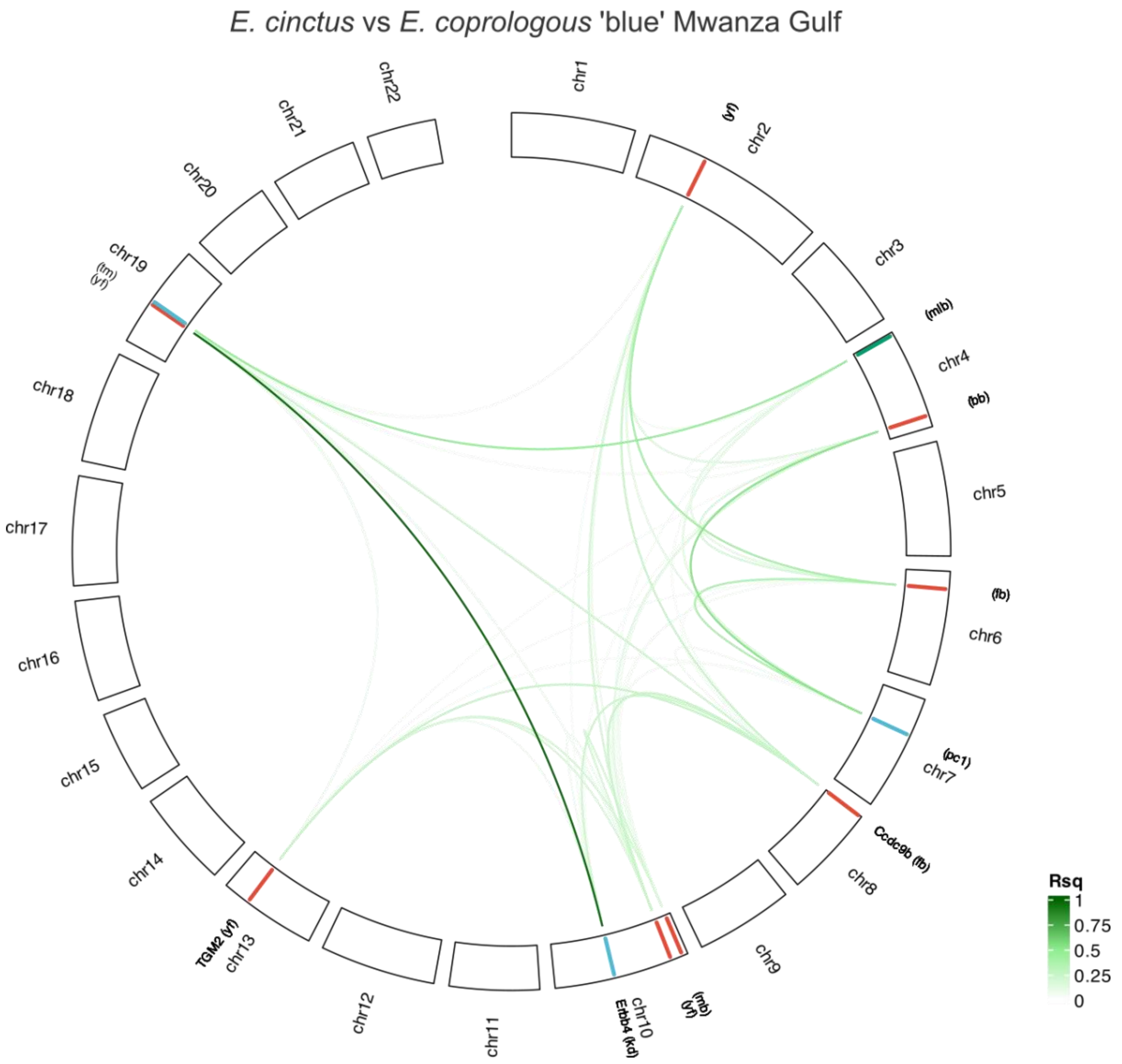
E. cinctus vs E. coprologous 'blue' Mwanza Gulf
chr1
(vf)
chr2
chr3
(mlb)
chr4
(bb)
chr5
(fb)
chr6
(pc1)
chr7
Ccdc9b (fb)
chr8
chr9
(mb)
(vf)
chr10
Erbb4 (kd)
chr11
chr12
TGM2 (vf)
chr13
chr14
chr15
chr16
chr17
chr18
chr19
(tm)
(vf)
chr20
chr21
chr22
Rsq
1
0.75
0.5
0.25
0

*Y. plumbus* vs *Y. pyrrocephalus* Makobe island

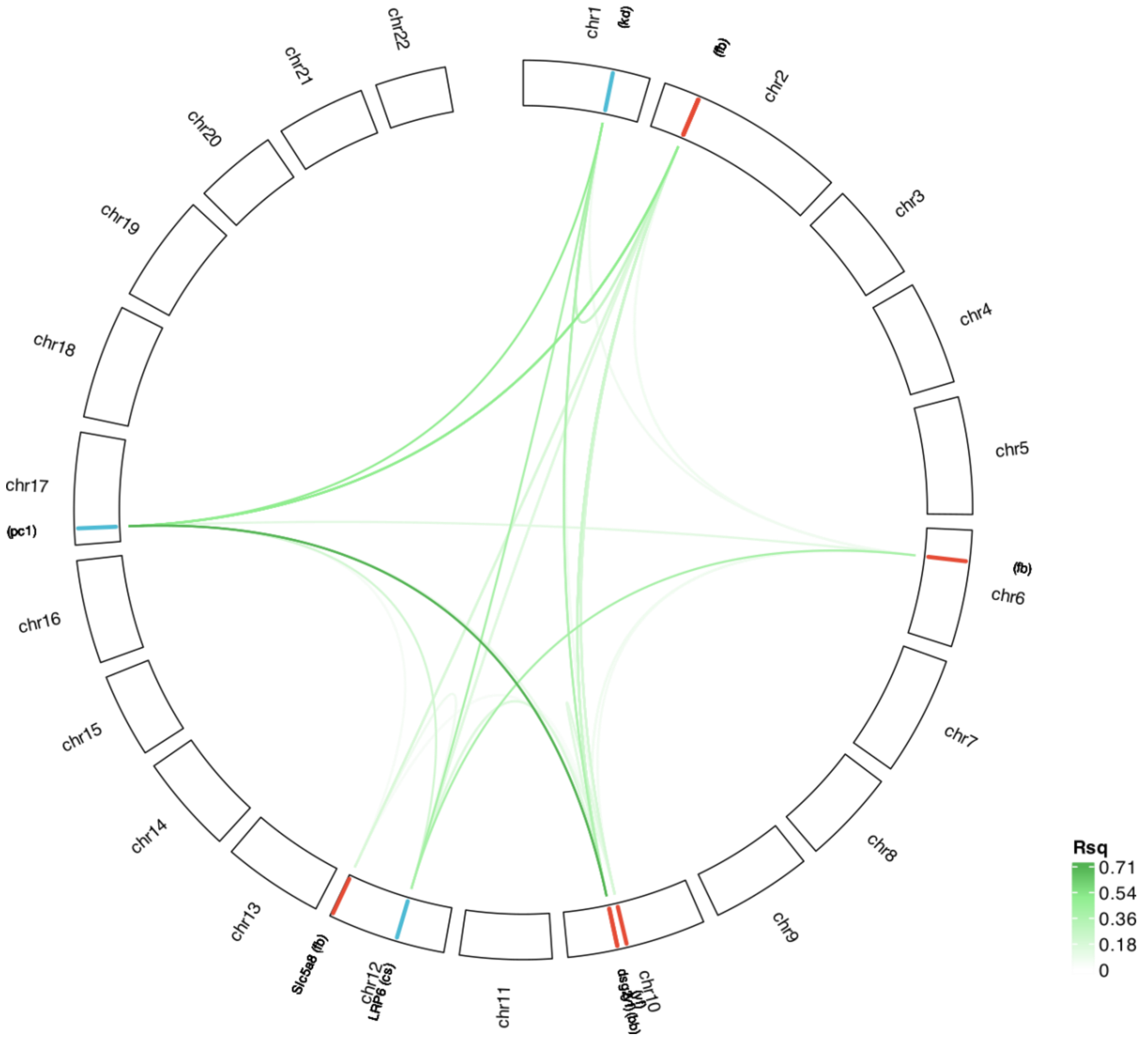

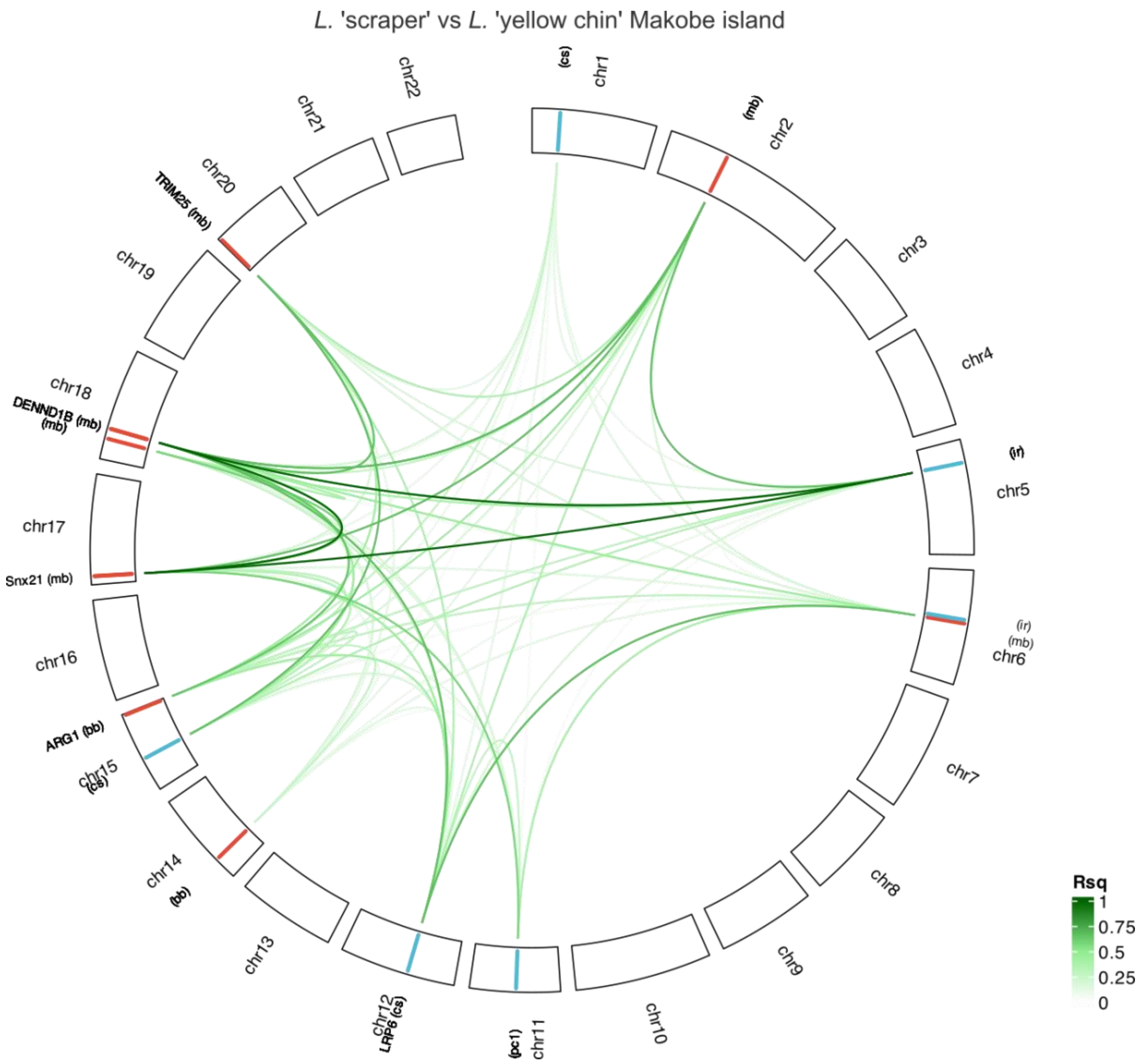
*L.* 'scraper' vs *L.* 'yellow chin' Makobe island
chr1
(cs)
chr2
(mb)
chr3
chr4
chr5
(ir)
chr6
(ir)
(mb)
chr7
chr8
chr9
chr10
chr11
(pc1)
chr12
LRP6 (cs)
chr13
chr14
(bb)
chr15
(cs)
ARG1 (bb)
chr16
chr17
Snx21 (mb)
chr18
DENND1B (mb)
(mb)
chr19
chr20
TRIM25 (mb)
chr21
chr22
Rsq
1
0.75
0.5
0.25
0

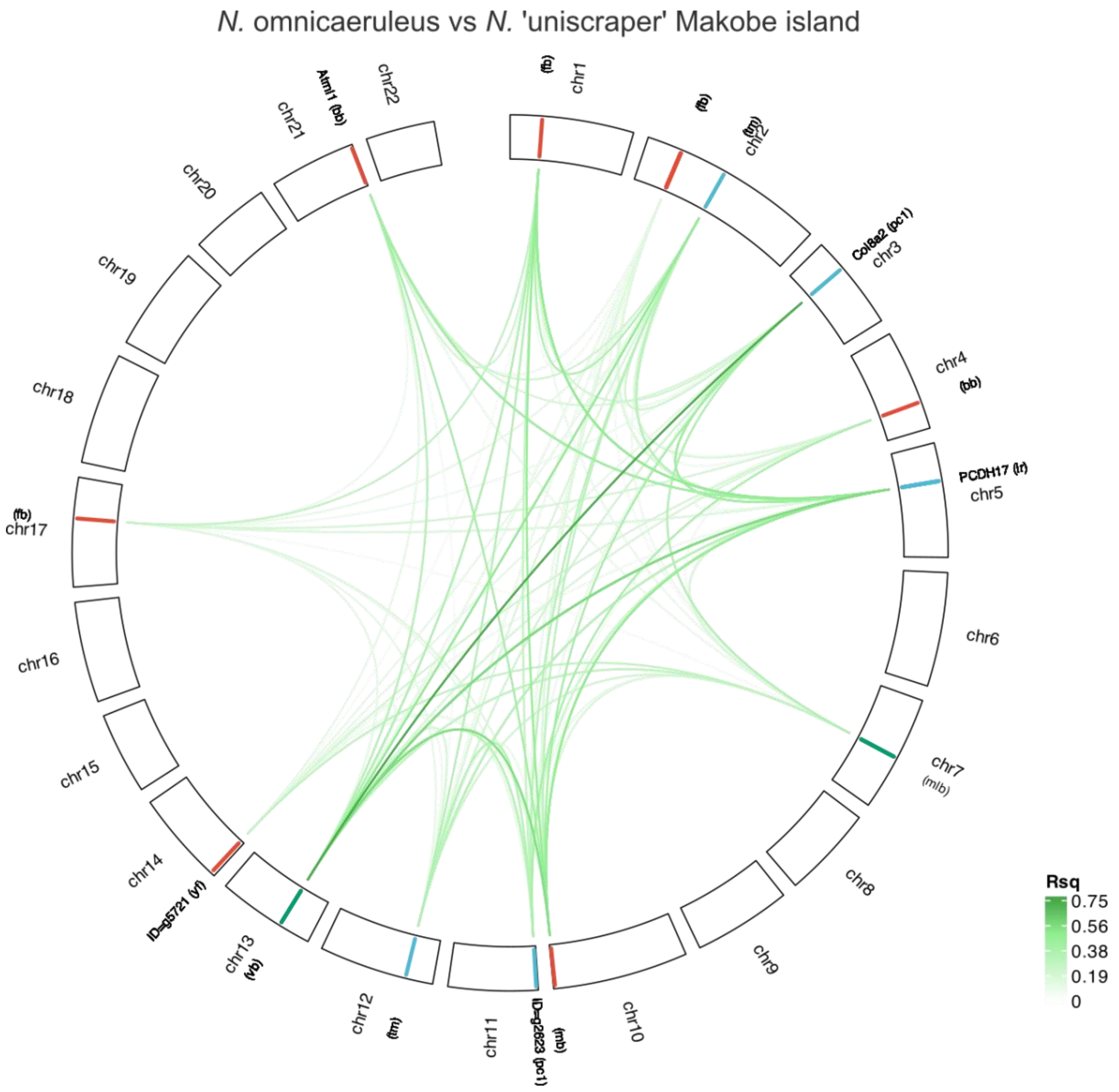
*N.* omnicaeruleus vs *N.* 'uniscraper' Makobe island
chr1
chr2
chr3
Col8a2 (pc1)
chr4
PCDH17 (lr)
chr5
chr6
chr7
(mlb)
chr8
chr9
chr10
chr11
ID=g2623 (pc1)
chr12
chr13
ID=g5721 (vf)
chr14
chr15
chr16
chr17
(fb)
chr18
chr19
chr20
chr21
Atml1 (bb)
chr22
Rsq
0.75
0.56
0.38
0.19
0

*N. gigas* vs *P. cyaneus* Makobe island

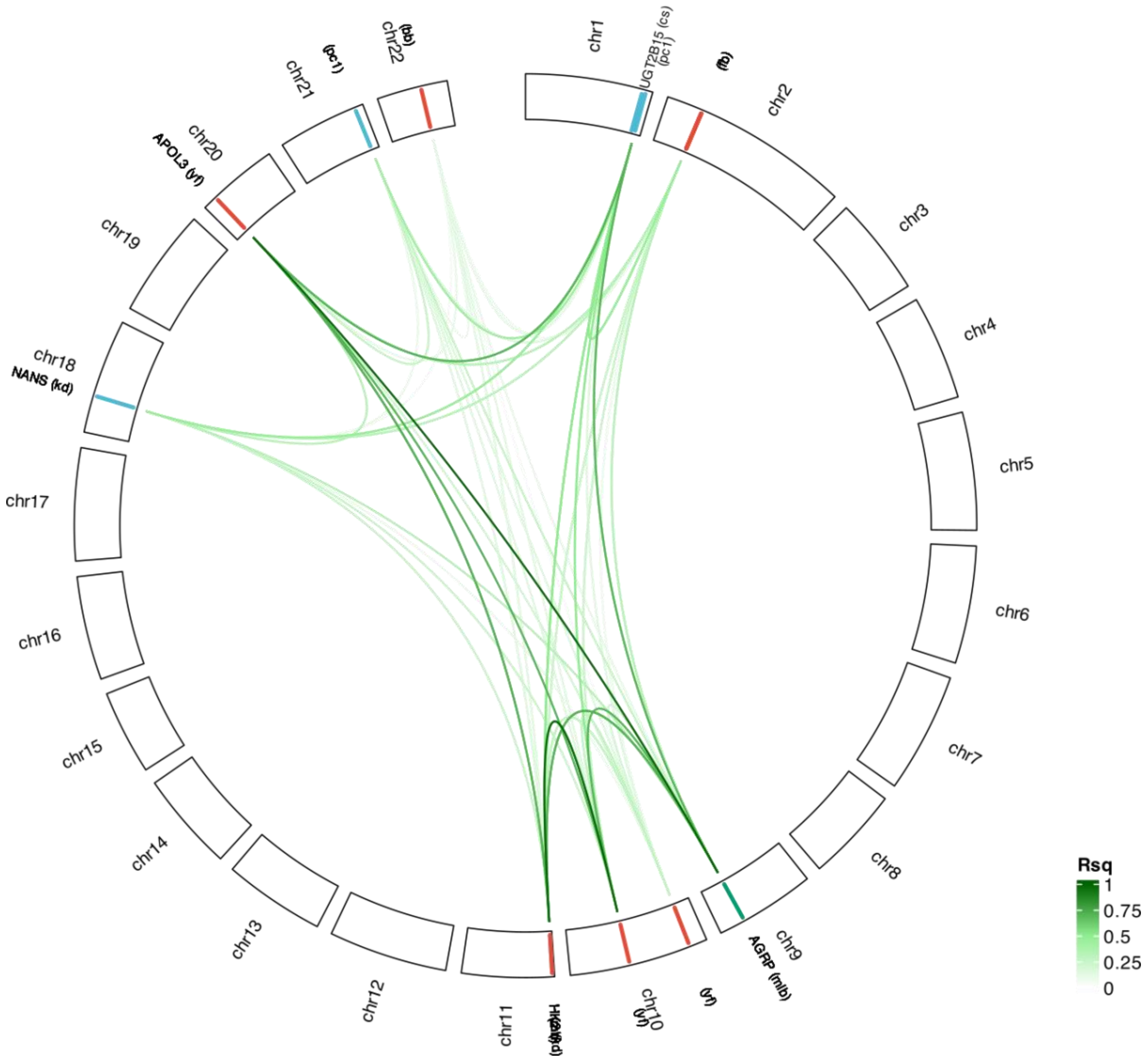

*M. mbipi* vs *P.* 'pink anal' Makobe island

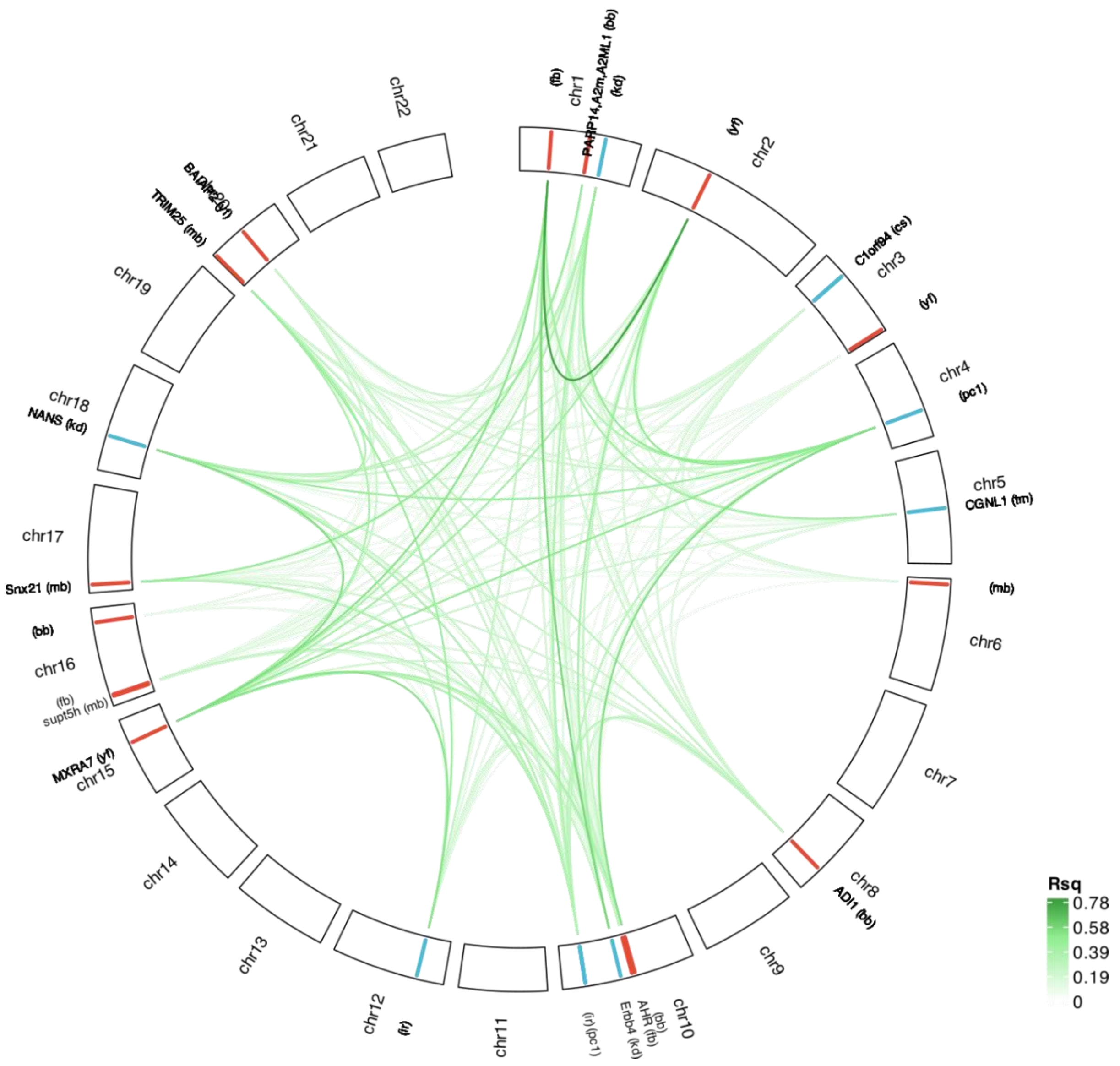

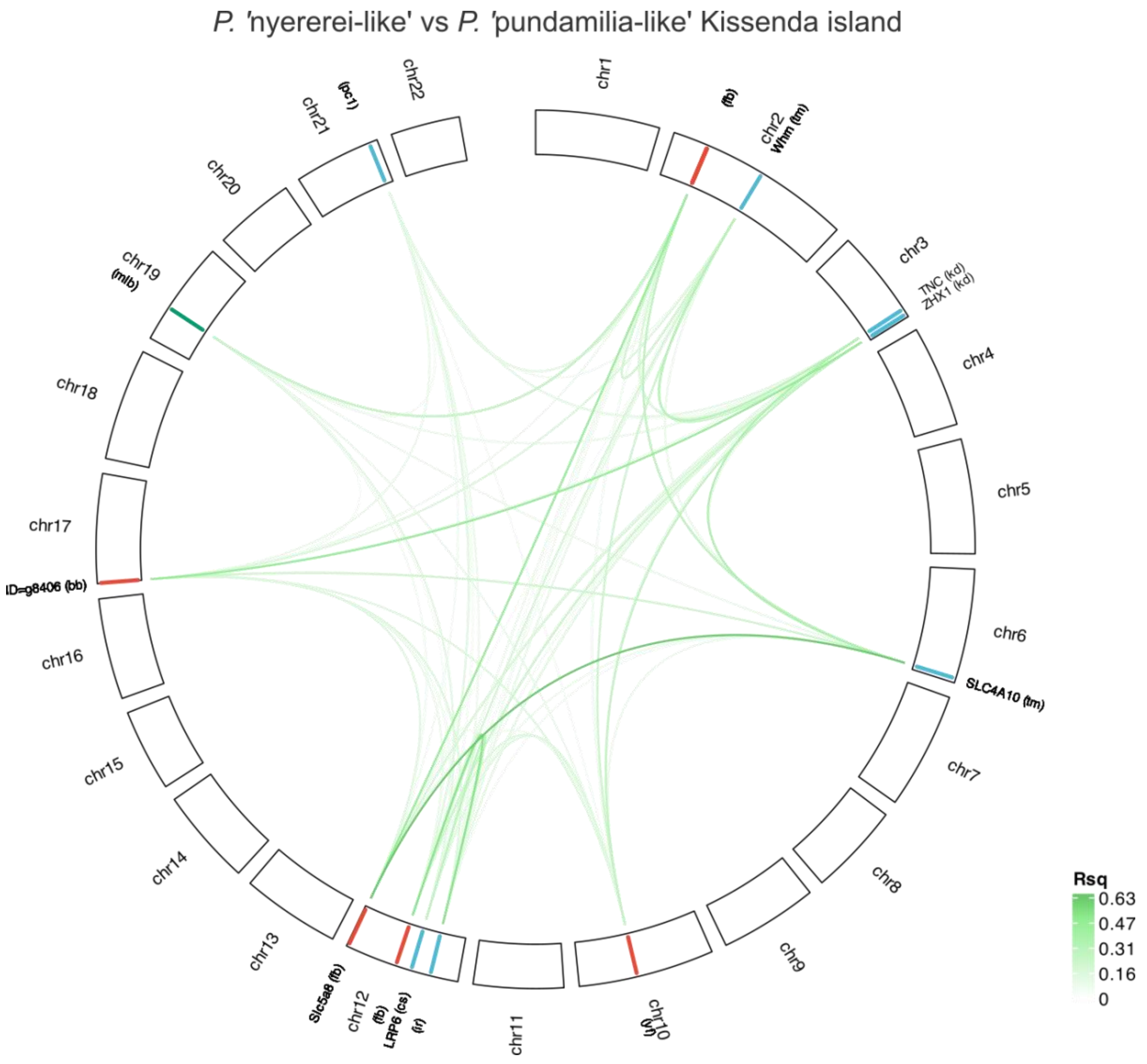

P. 'nyererei-like' vs P. 'pundamilia-like' Kissenda island
chr1
(fb)
chr2
Whm (tm)
chr3
TNC (kd)
ZHX1 (kd)
chr4
chr5
chr6
SLC4A10 (tm)
chr7
chr8
chr9
chr10
(yf)
chr11
chr12
(fb)
LRP6 (cs)
(rf)
Slc5a8 (fb)
chr13
chr14
chr15
chr16
ID=g8406 (bb)
chr17
chr18
chr19
(mlb)
chr20
chr21
(pc1)
chr22
Rsq
0.63
0.47
0.31
0.16
0

*P.* 'nyererei-like' vs *P.* 'pundamilia-like' Python island

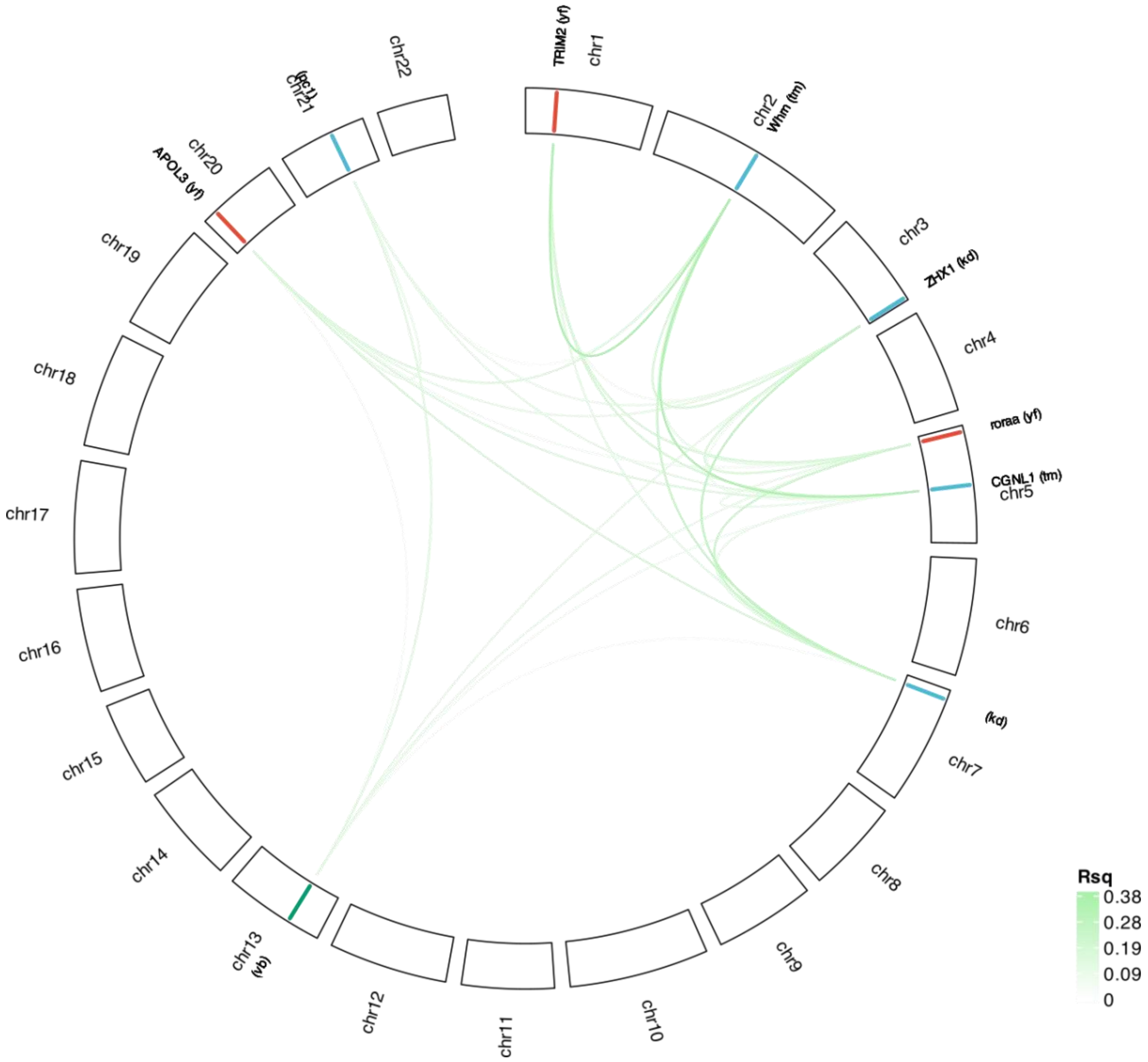

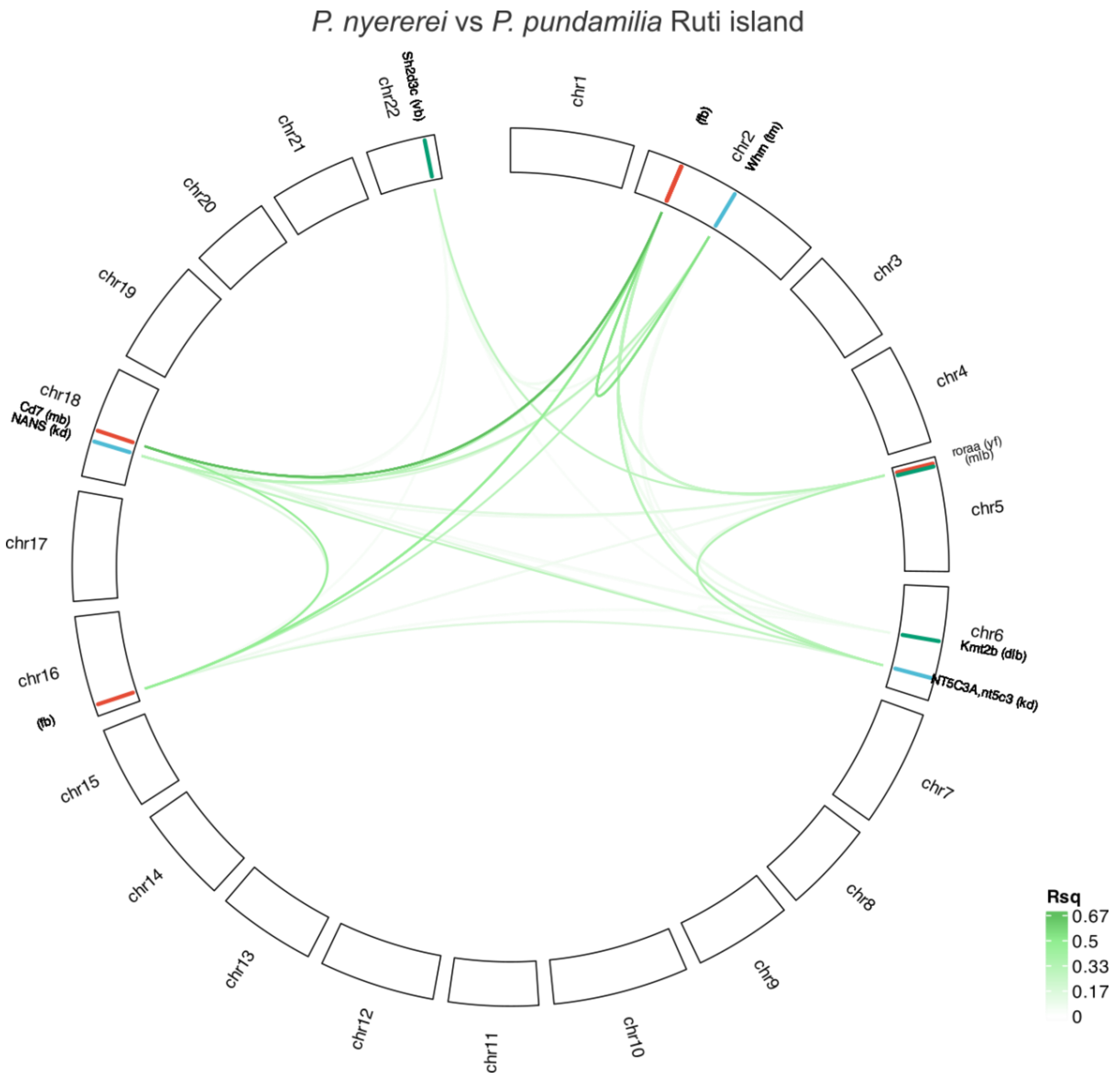

**Fig. S9 Combinatorial speciation in 10 sympatric cichlid sister-species in Lake Victoria through coupling of ecological and mating trait modules during combinatorial speciation.**
Continued from main Fig. 4. Genome-wide distribution of Linkage disequilibrium (LD; measured using $r^2$ statistic) coupled between sister species. Lines on the ideogram represent a top SNP and top 5% $F_{ST}$ and line colour denotes the trait complex the SNP was associated with in the RWAS: blue – ecomorphology, green-melanic stripe pattern, red-male nuptial colour as in Fig. 2. Trait abbreviations are printed in brackets next to the SNP ideogram. If SNP maps to gene, the gene name is shown. Higher LD ($r^2$) values are depicted by darker green colours. Only $r^2 > 0.01$ is shown. Only SNPs on different chromosomes and SNPs more than 100 Kbp away on the same chromosome were included in the analysis to exclude effects of physical linkage. Trait abbreviations are as follows: cusp shape (cs), and number of inner tooth rows in oral jaws (ir), lower pharyngeal jaw keel depth (kd) and molarisation of lower pharyngeal teeth (tm), as well are 3-dimensional (3D) craniofacial shape PC1

(pc1) and PC2 (pc2), vertical bars (vb), midlateral band (mlb) and dorsolateral band (dlb), flameback (fb), yellow (yf) or blue flank (bb), red chest (rc), and melanic body (mb).

#### Materials and Methods

##### Genome sequencing and assembly

DNA extraction was performed from blood of a freshly killed male *Pundamilia nyererei* individual with a phenol-chloroform protocol modified to limit degradation by DNAses. PacBio sequences were obtained through SMRT sequencing using 8 SMRT Cells on the Sequel System of Pacific BioScience. Simultaneously, paired-end Illumina sequences were generated from four closely related samples. The genome was assembled with PacBio sequences using flye v2.3.4 (*49*). Subsequently, genome polishing aimed to enhance the assembly's consensus sequence quality. One round of polishing with PacBio reads using arrow v2.2.2 (Pacific Bioscience) and two rounds with Illumina reads using pilon v1.22 (*50*) were applied. For scaffolding, additional mate-pair Illumina sequences previously obtained by Brawand et. al. (*51*) were downloaded from SRA (SRP004869) and quality trimmed using Trimmomatic v0.36 (*52*) (removing Illumina adapters, leading and trailing low quality bases (<3 or Ns), cutting the read when the average quality per base drops below 15 within a 4bp sliding window and removing reads shorter than 20bp). The trimmed mate-pair and paired-end Illumina reads were then used for scaffolding with SOAPdenovo2 v2.04.241 (*53*) and a k-mer size of 20. Scaffolds were ordered and oriented according to Feulner et al. [6] using their linkage map with ALLMAPS v0.8.4 (*54*). The final assembly underwent quality assessment with QUAST v4.6.0 (*55*) and BUSCO 3.0.2 (*56*). See Fig. S10 for bioinformatics pipeline.

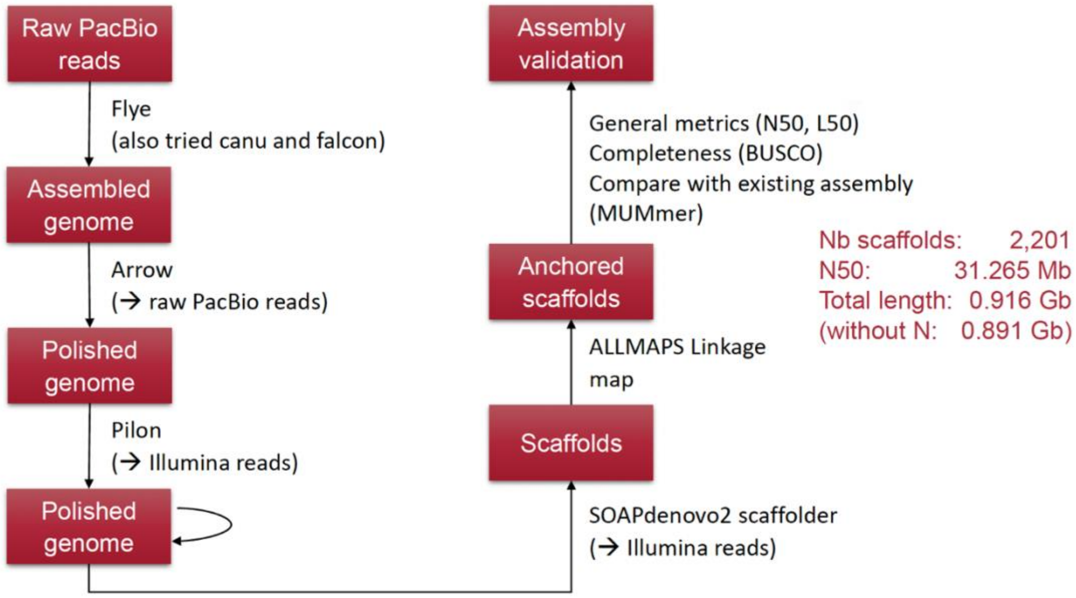

**Fig. S10 Genome assembly bioinformatics pipeline**

**Genome annotation**

Before annotation, repeats in the genome were masked using RepeatModeler2 v2.0.4 (*57*) and RepeatMasker v4.1.4 (*58*). PacBio HiFi IsoSeq reads were obtained for 5 different tissues (skin, brain, testits, eye, liver and kidney). The IsoSeq v3.8.2 pipeline from Pacific Bioscience was used to preprocess IsoSeq sequences and obtain transcript annotations on the assembled genome separately for each tissue. This includes lima to remove IsoSeq cDNA primers, isoseq3 refine to remove polyA tail and arifical concatemers, isoseq3 cluster to de novo isoform-level clustering, pbmm2 align isoform-level clusters to the genomes and isoseq3 collapse to collapse redundant transcripts based on exonic structures. Predicted protein-coding regions in the annotated transcripts were identified using GeneMarkS-T (*59*), and AGAT v1.0.0 (*60*) corrected the resulting gtf file. Additionally, RNAseq data from Brawand et al. (*51*) were downloaded from SRA (SRP010906), quality trimmed using cutadapt v3.4.1 [14] (removing Illumina adapters and reads shorter than 20bp) and mapped to the genome with HiSat2 v2.2.1 (*61*). The BRAKER3 annotation pipeline v3.0.2 (*62*) was employed to annotate the repeat-masked genome using the mapped RNAseq data and vertebrate protein data from OrthoDB v11 (*63*). The annotation predictions from BRAKER3 and IsoSeq data were combined using TSEBRA (*64*) with the "long_reads_filtered" configuration file.

To ensure a comprehensive annotation, available IsoSeq data from three missing tissues (PJ, eye, and blue skin) from the sister species *Pundamilia pundamilia* were incorporated. Transcript annotation from this data followed the same procedure as previously described. The new annotations were then merged with the combined annotation using TSEBRA with the "long_reads_filtered" configuration file, along with the "keep_gtf" option to ensure inclusion of all transcripts from the combined annotation file.Gffcompare v0.12.6 (*65*) was used to identify complete matches between individual IsoSeq annotations and the merged annotation in order to add tissue and species information to the final annotation file. Functional information was incorporated through ncbi-blastp v2.10.1+ (*66*) searches of coding amino acid sequences against the UniProt/Swiss-Prot database (*67*) with an e-value threshold of 10 and InterProScan v5.51.85.0 (*68*) with mapping to all default databases and gene ontology terms. AGAT was used to combine and integrate these functional annotations into the annotation file with a blast e-value threshold of 0.01. Finally, Ensemble IDs were obtained from Ensemble BioMart (*69*). See Fig. S11 for bioinformatics pipeline.

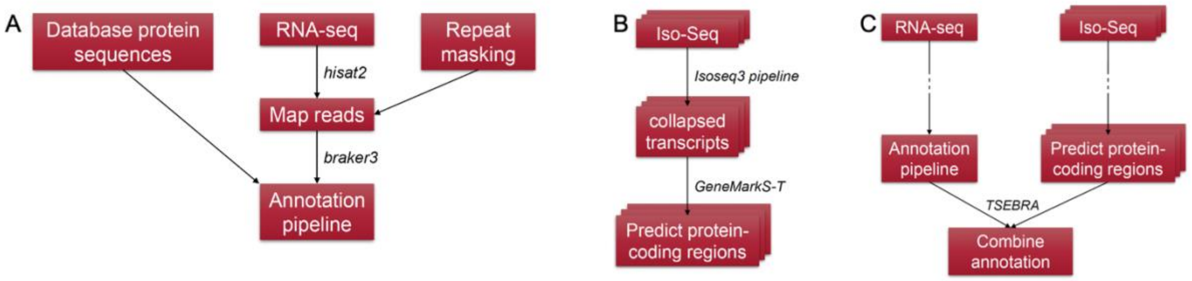

**Fig. S11 Genome annotation bioinformatics pipeline used to assemble the *P. nyererei* v3 genome**

**Phenotyping and geometric-morphometric analysis**

Male colouration (yellow flank, flameback, red chest, blue body, melanic body), melanic body patterning (vertical bars, midlateral band, dorsolateral band), and dentition (oral teeth cusp shape, number of inner row of upper oral teeth, pharyngeal jaw keel depth, pharyngeal jaw molarisation) traits were phenotyped across 107 species (File S1) based on fish images and data from (*70*) as present/absent or assigned to categories (Fig. S2) that can be visualised across the phylogeny in Fig. 1. To determine morphological variation in craniofacial shape, we CT scanned 318 individuals from 105 species using a SkyScan 1272 and a SkyScan 2214 CT scanners at UniBern and a Bruker Skyscan 1273 at Rice University. These scans were rendered into 3D objects in the open-source software 3D-Slicer and landmarked using the Markups module. We included 25 landmarks on each specimen, covering half of the head (Fig. S1a). These landmarks were then analysed in the Geomorph package in R using a Procrustes Superimposition and Generalized Principal Components Analysis. An analysis of morphological disparity (morphol.disparity) was performed on species averages of shape to assess the amount of morphological variation across diet category and genus. Principal components 1 and 2 from this analysis were used a craniofacial shape trait for the genotype-phenotype association analysis (Fig. S1, File S1).

**Alignment and SNP calling**

Whole genome re-sequencing data of 324 individuals from (*71–73*) was re-analysed for this manuscript. Paired 150bp reads for all individuals was mapped to the improved *P. nyererei* v3.0 reference using bwa-mem (*74*) (see File S1 for mapping statistics). Duplicates were marked using samtools and SNP calling was conducted using bcftools (*75*). SNPs were filtered using bcftools (*75*) (per sample genotype depth = 20x, Phred quality= 30, <1% missingness and minor allele frequency > 0.05. Missing genotypes were imputed using Beagle (*76*) before genotype-phenotype association analysis.

**RWAS: radiation wide genotype-phenotype association analysis**

We implemented the Bayesian sparse linear mixed model (BSLMM) in GEMMA (*77*) that integrates features of standard linear mixed models - assuming that all genetic variants contribute small effects to phenotype (polygenic) - and sparse regression models, which assumed that only a small subset of variants have non-zero affect phenotype (oligo/mono-genic). Population structure was accounted for by estimating a kinship relatedness matrix as a random effect in the model. Sex was used as a covariate as it is especially important for male nuptial colouration phenotypes. BSLMM utilises Markov chain Monte Carlo (MCMC) to estimate several key parameters: the proportion of phenotypic variation explained by each SNP included in the analysis (PVE), the proportion explained by SNPs of large effect (PGE), which are those with a non-zero effect on phenotype, and the number of large-effect SNPs needed to explain PGE (nSNPs). For each SNP, GEMMA computes an effect size coefficient (β) and a posterior inclusion probability (PIP). A non-zero value of β in one MCMC iteration indicates that the marker influences phenotypic variation. As β can be positive or negative depending on the direction of association, absolute values are presented for clarity. PIP represents the proportion of MCMC iterations in which a SNP is estimated to have a non-zero effect on phenotype ($\beta \neq 0$). As all SNPs are treated simultaneously in a single model, correction for multiple testing is not required. To ensure robust results, we conducted 10 independent iterations of the BSLMM (following (*78*)), each consisting of 20 million MCMC steps with a burn-in of 5 million steps. For each SNP, we calculated the median β and PIP values across the 10 independent MCMC chains. Across these independent runs, consistent results emerged, pinpointing the strongest SNP associations. For downstream analysis we applied top 99.99$^{th}$ percentile & PIP > 0.01 threshold for binary/categorical traits and top 99.9$^{th}$ percentile & PIP > 0.01 threshold for continuous traits; denoted top SNPs with potentially meaningful association based on (*78*).

**Machine learning prediction of genotype-phenotype association results for binary traits**

To evaluate the predictive accuracy of candidate loci identified by the association analyses, we employed a machine learning framework using Random Forest (RF) classification for eight binary body colour and patterning traits. Analyses were restricted to as male nuptial colour is not expressed in females. The genotype matrix, filtered for top RWAS SNPs, was merged with phenotype data. For each trait, we implemented a five-fold cross-validation procedure. Within each fold, RF classifiers were trained on four-fifths of the data and evaluated on the held-out fold using 1,500 trees. Model performance was quantified using Receiver Operating Characteristic (ROC) curves. SNP importance was extracted using Mean Decrease Accuracy (MDA) and Mean Decrease Gini (MDG). To identify robust genomic drivers, we implemented a “Random Forest Fishing” (RFF) procedure (*79*) within each cross-validation fold. Briefly, an initial bait set consisting of the top five SNPs ranked by PIP was iteratively combined with random pools of candidate SNPs. At each iteration, a RF model was fitted, and SNPs were ranked by MDA; the top 50% were retained to update the bait set. This process was repeated for multiple iterations, yielding a final set of selected SNPs per fold. To quantify selection stability, we calculated SNP consistency as the proportion of cross-validation folds in which a SNP was retained by the RFF procedure (range: 0–1). Finally, we assessed concordance between GWAS results and machine learning importance metrics by testing for associations between SNP PIP scores and RF importance measures (MDA) using Pearson correlations and linear regression. All statistical analyses and visualisations were performed in R (v4.3) using the randomForest, pROC, data.table, dplyr, ggplot2, patchwork, and ggrepel packages.

**Phylogenomics**

To infer the evolutionary relationships among individuals, we generated a high-quality, linkage-reduced SNP alignment from whole-genome resequencing data. Variant sites were first filtered to retain only informative positions using bcftools (*75*) by requiring at least one homozygous reference (0/0) and one homozygous alternate (1/1) genotype per site. This removed invariant loci that do not contribute phylogenetic signal. To reduce linkage and computational burden, the filtered SNP set was thinned to retain only one SNP per 1 kb using VCFtools (*80*), ensuring an approximately independent SNP set suitable for phylogenetic inference. The thinned VCF was converted to PHYLIP format using the vcf2phylip pipeline (*81*), producing a multi-sequence alignment containing only biallelic, informative, and widely spaced SNPs. Phylogenetic reconstruction was performed using IQ-TREE v2 (*82*), employing the GTR+G substitution model. Node support was evaluated using 1,000 ultrafast bootstrap replicates and 1,000 SH-aLRT tests.

**GO enrichment analysis**

Gene set enrichment analysis was conducted using topGO (v2.36.0)(*83*) using the method weight to account for GO topology and Fisher's exact test to correct for multiple testing for the enrichment analysis. We used GO annotations from our *P. nyererei* v3 annotation.

**FST and Linkage Disequilibrium calculation**

Pairwise FST was calculated in 20kb windows non-overlapping windows between sister species (minimum n=3 per species) using vcftools (*80*). We calculated Linkage disequilibrium (LD), which is the non-random association of alleles at two loci, using the $r^2$ statistic (*84*) computed as:

$$r^2 = \frac{(D^2)}{(pA(1-pA)) * (pB(1-pB))}$$

Where $D$ is the linkage disequilibrium coefficient, calculated as $D = pAB - pApB$ , where $pAB$ is the frequency of the haplotype containing alleles $A$ and $B$ and $pA$ and $pB$ are the allele frequencies for the individual loci. $r^2$ statistic of 1 indicates complete LD (the alleles are always inherited together) and a value of 0 indicates no LD (the alleles are inherited independently). Only SNPs on different chromosomes and SNPs more than 100 Kbp away on the same chromosome were included in the analysis to exclude effects of physical linkage in all LD analyses.

**Supplementary Files**

**File S1 A excel sheet of all phenotypes and associated meta data of samples; top RWAS SNPs with associated information, and per sample coverage.**

51. D. Brawand, C. E. Wagner, Y. I. Li, M. Malinsky, I. Keller, S. Fan, O. Simakov, A. Y. Ng, Z. W. Lim, E. Bezault, J. Turner-Maier, J. Johnson, R. Alcazar, H. J. Noh, P. Russell, B. Aken, J. Alföldi, C. Amemiya, N. Azzouzi, J.-F. Baroiller, F. Barloy-Hubler, A. Berlin, R. Bloomquist, K. L. Carleton, M. a. Conte, H. D'Cotta, O. Eshel, L. Gaffney, F. Galibert, H. F. Gante, S. Gnerre, L. Greuter, R. Guyon, N. S. Haddad, W. Haerty, R. M. Harris, H. a. Hofmann, T. Hourlier, G. Hulata, D. B. Jaffe, M. Lara, A. P. Lee, I. MacCallum, S. Mwaiko, M. Nikaido, H. Nishihara, C. Ozouf-Costaz, D. J. Penman, D. Przybylski, M. Rakotomanga, S. C. P. Renn, F. J. Ribeiro, M. Ron, W. Salzburger, L. Sanchez-Pulido, M. E. Santos, S. Searle, T. Sharpe, R. Swofford, F. J. Tan, L. Williams, S. Young, S. Yin, N. Okada, T. D. Kocher, E. a. Miska, E. S. Lander, B. Venkatesh, R. D. Fernald, A. Meyer, C. P. Ponting, J. T. Streelman, K. Lindblad-

Toh, O. Seehausen, F. Di Palma, The genomic substrate for adaptive radiation in African cichlid fish. *Nature* **513**, 375–381 (2014).

52. A. M. Bolger, M. Lohse, B. Usadel, Trimmomatic: A flexible trimmer for Illumina sequence data. *Bioinformatics* **30**, 2114–2120 (2014).
53. R. Luo, B. Liu, Y. Xie, Z. Li, W. Huang, J. Yuan, G. He, Y. Chen, Q. Pan, Y. Liu, J. Tang, G. Wu, H. Zhang, Y. Shi, Y. Liu, C. Yu, B. Wang, Y. Lu, C. Han, D. W. Cheung, S. M. Yiu, S. Peng, Z. Xiaoqian, G. Liu, X. Liao, Y. Li, H. Yang, J. Wang, T. W. Lam, J. Wang, SOAPdenovo2: An empirically improved memory-efficient short-read de novo assembler. *Gigascience* **1**, 1–6 (2012).
54. H. Tang, X. Zhang, C. Miao, J. Zhang, R. Ming, J. C. Schnable, P. S. Schnable, E. Lyons, J. Lu, ALLMAPS: Robust scaffold ordering based on multiple maps. *Genome Biol.* **16**, 1–15 (2015).
55. A. Mikheenko, V. Saveliev, A. Gurevich, MetaQUAST: evaluation of metagenome assemblies. *Bioinformatics* **32**, 1088–1090 (2016).
56. R. M. Waterhouse, M. Seppey, F. A. Simao, M. Manni, P. Ioannidis, G. Klioutchnikov, E. V. Kriventseva, E. M. Zdobnov, BUSCO Applications from Quality Assessments to Gene Prediction and Phylogenomics. *Mol. Biol. Evol.* **35**, 543–548 (2018).
57. A. Smit, R. Hubley, RepeatModeler. Open-1.0 [Preprint] (2015). http://www.repeatmasker.org.
58. A. Smit, R. Hubley, P. Green, RepeatMasker. Open-4.0 [Preprint] (2015). http://www.repeatmasker.org.
59. S. Tang, A. Lomsadze, M. Borodovsky, Identification of protein coding regions in RNA transcripts. *Nucleic Acids Res.* **43**, e78–e78 (2015).
60. J. Dainat, AGAT: Another Gff Analysis Toolkit to handle annotations in any GTF/GFF format. Zenodo v1.0 [Preprint].
61. D. Kim, B. Langmead, S. L. Salzberg, HISAT: A fast spliced aligner with low memory requirements. *Nat. Methods* **12**, 357–360 (2015).
62. T. Brůna, K. J. Hoff, A. Lomsadze, M. Stanke, M. Borodovsky, BRAKER2: automatic eukaryotic genome annotation with GeneMark-EP+ and AUGUSTUS supported by a protein database. *NAR Genom. Bioinform.* **3**, lqaa108 (2021).
63. D. Kuznetsov, F. Tegenfeldt, M. Manni, M. Seppey, M. Berkeley, E. V. Kriventseva, E. M. Zdobnov, OrthoDB v11: annotation of orthologs in the widest sampling of organismal diversity. *Nucleic Acids Res.* **51**, D445–D451 (2023).
64. L. Gabriel, K. J. Hoff, T. Brůna, M. Borodovsky, M. Stanke, TSEBRA: transcript selector for BRAKER. *BMC Bioinformatics* **22**, 1–12 (2021).
65. G. Pertea, M. Pertea, GFF Utilities: GffRead and GffCompare [version 1; peer review: 3 approved] . *F1000Res.* **9** (2020).

66. C. Camacho, G. Coulouris, V. Avagyan, N. Ma, J. Papadopoulos, K. Bealer, T. L. Madden, BLAST+: Architecture and applications. *BMC Bioinformatics* **10** (2009).
67. E. Coudert, S. Gehant, E. de Castro, M. Pozzato, D. Baratin, T. Neto, C. J. A. Sigrist, N. Redaschi, A. Bridge, T. U. Consortium, A. J. Bridge, L. Aimo, G. Argoud-Puy, A. H. Auchincloss, K. B. Axelsen, P. Bansal, D. Baratin, T. M. B. Neto, M.-C. Blatter, J. T. Bolleman, E. Boutet, L. Breuza, B. C. Gil, C. Casals-Casas, K. C. Echioukh, E. Coudert, B. Cuche, E. de Castro, A. Estreicher, M. L. Famiglietti, M. Feuermann, E. Gasteiger, P. Gaudet, S. Gehant, V. Gerritsen, A. Gos, N. Gruaz, C. Hulo, N. Hyka-Nouspikel, F. Jungo, A. Kerhornou, P. Le Mercier, D. Lieberherr, P. Masson, A. Morgat, V. Muthukrishnan, S. Paesano, I. Pedruzzi, S. Pilbout, L. Pourcel, S. Poux, M. Pozzato, M. Pruess, N. Redaschi, C. Rivoire, C. J. A. Sigrist, K. Sonesson, S. Sundaram, A. Bateman, M.-J. Martin, S. Orchard, M. Magrane, S. Ahmad, E. Alpi, E. H. Bowler-Barnett, R. Britto, H. B.- A-Jee, A. Cukura, P. Denny, T. Dogan, T. Ebenezer, J. Fan, P. Garmiri, L. J. da Costa Gonzales, E. Hatton-Ellis, A. Hussein, A. Ignatchenko, G. Insana, R. Ishtiaq, V. Joshi, D. Jyothi, S. Kandasaamy, A. Lock, A. Luciani, M. Lugaric, J. Luo, Y. Lussi, A. MacDougall, F. Madeira, M. Mahmoudy, A. Mishra, K. Moulang, A. Nightingale, S. Pundir, G. Qi, S. Raj, P. Raposo, D. L. Rice, R. Saidi, R. Santos, E. Speretta, J. Stephenson, P. Totoo, E. Turner, N. Tyagi, P. Vasudev, K. Warner, X. Watkins, R. Zaru, H. Zellner, C. H. Wu, C. N. Arighi, L. Arminski, C. Chen, Y. Chen, H. Huang, K. Laiho, P. McGarvey, D. A. Natale, K. Ross, C. R. Vinayaka, Q. Wang, Y. Wang, Annotation of biologically relevant ligands in UniProtKB using ChEBI. *Bioinformatics* **39** (2023).
68. P. Jones, D. Binns, H. Y. Chang, M. Fraser, W. Li, C. McAnulla, H. McWilliam, J. Maslen, A. Mitchell, G. Nuka, S. Pesseat, A. F. Quinn, A. Sangrador-Vegas, M. Scheremetjew, S. Y. Yong, R. Lopez, S. Hunter, InterProScan 5: Genome-scale protein function classification. *Bioinformatics* **30**, 1236–1240 (2014).
69. F. J. Martin, M. R. Amode, A. Aneja, O. Austine-Orimoloye, A. G. Azov, I. Barnes, A. Becker, R. Bennett, A. Berry, J. Bhai, S. K. Bhurji, A. Bignell, S. Boddu, P. R. Branco Lins, L. Brooks, S. B. Ramaraju, M. Charkhchi, A. Cockburn, L. Da Rin Fiorretto, C. Davidson, K. Dodiya, S. Donaldson, B. El Houdaigui, T. El Naboulsi, R. Fatima, C. G. Giron, T. Genez, G. S. Ghattaoraya, J. G. Martinez, C. Guijarro, M. Hardy, Z. Hollis, T. Hourlier, T. Hunt, M. Kay, V. Kaykala, T. Le, D. Lemos, D. Marques-Coelho, J. C. Marugán, G. A. Merino, L. P. Mirabueno, A. Mushtaq, S. N. Hossain, D. N. Ogeh, M. P. Sakthivel, A. Parker, M. Perry, I. Piliota, I. Prosovetskaia, J. G. Perez-Silva, A. I. A. Salam, N. Saraiva-Agostinho, H. Schuilenburg, D. Sheppard, S. Sinha, B. Sipos, W. Stark, E. Steed, R. Sukumaran, D. Sumathipala, M. M. Suner, L. Surapaneni, K. Sutinen, M. Szpak, F. F. Tricomi, D. Urbina-Gómez, A. Veidenberg, T. A. Walsh, B. Walts, E. Wass, N. Willhoft, J. Allen, J. Alvarez-Jarreta, M. Chakiachvili, B. Flint, S. Giorgetti, L. Haggerty, G. R. Ilsley, J. E. Loveland, B.

Moore, J. M. Mudge, J. Tate, D. Thybert, S. J. Trevanion, A. Winterbottom, A. Frankish, S. E. Hunt, M. Ruffier, F. Cunningham, S. Dyer, R. D. Finn, K. L. Howe, P. W. Harrison, A. D. Yates, P. Flicek, Ensembl 2023. *Nucleic Acids Res.* **51**, D933–D941 (2023).

70. O. Seehausen, *Lake Victoria Rock Cichlids* (Verduyn Cichlids, 1996).
71. M. D. McGee, S. R. Borstein, J. I. Meier, D. A. Marques, S. Mwaiko, A. Taabu, M. A. Kishe, B. O'Meara, R. Bruggmann, L. Excoffier, O. Seehausen, The ecological and genomic basis of explosive adaptive radiation. *Nature* **586**, 75–79 (2020).
72. J. I. Meier, M. D. McGee, D. A. Marques, S. Mwaiko, M. Kishe, S. Wandera, D. Neumann, H. Mrosso, L. J. Chapman, C. A. Chapman, L. Kaufman, A. Taabu-Munyaho, C. E. Wagner, R. Bruggmann, L. Excoffier, O. Seehausen, Cycles of fusion and fission enabled rapid parallel adaptive radiations in African cichlids. *Science (1979).* **381** (2023).
73. D. A. Marques, J. I. Meier, M. D. McGee, M. P. Haesler, S. Mwaiko, M. A. Kishe, A. Taabu-Munyaho, L. Chapman, C. A. Chapman, S. B. Wandera, C. E. Wagner, L. E. Excoffier, O. Seehausen, Genomes reveal age and demographic consequence of ultrafast adaptive radiation. *bioRxiv*, 2025.03.07.638630 (2025).
74. H. Li, Aligning sequence reads, clone sequences and assembly contigs with BWA-MEM. (2013).
75. P. Danecek, J. K. Bonfield, J. Liddle, J. Marshall, V. Ohan, M. O. Pollard, A. Whitwham, T. Keane, S. A. McCarthy, R. M. Davies, H. Li, Twelve years of SAMtools and BCFtools. *Gigascience* **10** (2021).
76. B. L. Browning, Beagle 3.3.2. 1–30 (2011).
77. X. Zhou, P. Carbonetto, M. Stephens, Polygenic Modeling with Bayesian Sparse Linear Mixed Models. *PLoS Genet.* **9** (2013).
78. A. A. Comeault, V. Soria-Carrasco, Z. Gompert, T. E. Farkas, C. A. Buerkle, T. L. Parchman, P. Nosil, Genome-wide association mapping of phenotypic traits subject to a range of intensities of natural selection in Timema cristinae. *American Naturalist* **183**, 711–727 (2014).
79. W. Yang, C. Charles Gu, Random forest fishing: a novel approach to identifying organic group of risk factors in genome-wide association studies. *European Journal of Human Genetics 2014 22:2* **22**, 254–259 (2013).
80. P. Danecek, A. Auton, G. Abecasis, C. A. Albers, E. Banks, M. A. DePristo, R. E. Handsaker, G. Lunter, G. T. Marth, S. T. Sherry, G. McVean, R. Durbin, The variant call format and VCFtools. *Bioinformatics* **27**, 2156–2158 (2011).
81. E. M. Ortiz, vcf2phylip v2.0: convert a VCF matrix into several matrix formats for phylogenetic analysis. [Preprint] (2019). https://doi.org/10.5281/ZENODO.2540861.
82. B. Q. Minh, H. A. Schmidt, O. Chernomor, D. Schrempf, M. D. Woodhams, A. Von Haeseler, R. Lanfear, E. Teeling, IQ-TREE 2: New Models and Efficient Methods for Phylogenetic Inference in the Genomic Era. *Mol. Biol. Evol.* **37**, 1530–1534 (2020).

83. A. Alexa, J. Rahnenfuhrer, topGO: Enrichment analysis for gene ontology. R package version 2.38.1. [Preprint] (2019).
84. W. G. Hill, A. Robertson, Linkage disequilibrium in finite populations. *Theoretical and Applied Genetics* **38**, 226–231 (1968).
